# The *Drosophila* Individual Activity Monitoring and Detection System (DIAMonDS)

**DOI:** 10.1101/2020.06.05.136085

**Authors:** Ki-Hyeon Seong, Taishi Matsumura, Yuko Shimada-Niwa, Ryusuke Niwa, Siu Kang

**Affiliations:** RIKEN Cluster for Pioneering Research, RIKEN Tsukuba Institute, 3-1-1 Koyadai, Tsukuba, Ibaraki 305-0074, Japan; Graduate School of Science and Engineering, Yamagata University, Jonan, Yonezawa, Yamagata 992-8510, Japan; AMED-CREST, AMED, 1-7-1 Otemachi, Chiyoda-ku, Tokyo 100-0004, Japan; Life Science Center for Survival Dynamics, University of Tsukuba, Tennoudai 1-1-1, Tsukuba 305-8577, Japan

## Abstract

Here, we have developed DIAMonDS (*Drosophila* Individual Activity Monitoring and Detection System) comprising time-lapse imaging by a charge-coupled device (CCD) flatbed scanner and Sapphire, a novel algorithm and web application. DIAMonDS automatically and sequentially identified the transition time points of multiple life cycle events such as pupariation, eclosion, and death in individual flies at high temporal resolution and on a large scale. DIAMonDS performed simultaneous multiple scans to measure individual deaths (≤ 1,152 flies per scanner) and pupariation and eclosion timings (≤ 288 flies per scanner) under various chemical exposures, environmental conditions, and genetic backgrounds. DIAMonDS correctly identified 74–85% of the pupariation and eclosion events and ∼92% of the death events within ±10 scanning frames. This system is a powerful tool for studying the influences of genetic and environmental factors on fruit flies and efficient, high-throughput genetic and chemical screening in drug discovery.

## Introduction

Animal life develops through a sequence of characteristic events and stages. Embryogenesis starts with the fertilization of an egg. Thence, the embryo transitions to the juvenile stage at a certain time point. This stage persists for a certain period, during which the animal grows and develops into the adult stage in which it reproduces, senesces, and dies. The length of each stage and the timing of each developmental transition are determined by complex interactions between genetic and environmental factors. Thus, developmental timing may be defined as a phenotype. Accurate determination of developmental timing helps elucidate the molecular and physiological bases of biological events and may uncover new factors regulating development and lifespan.

Nevertheless, it is difficult to establish the precise time points of the developmental transitions in many animals including humans. One reason is that there may be no clear boundaries between stages throughout the entire lifetime of the organism. Consequently, it has been impractical to use developmental timing as a phenotype for investigations in developmental biology. However, holometabolous insects such as the fruit fly *Drosophila melanogaster* are important exceptions to this rule. These animals pass through four life stages, namely embryo, larva, pupa, and adult. Transitions between these stages are accompanied by drastic morphological events and behavioral changes including hatching, pupariation, eclosion, and death. Therefore, the time point of each developmental transition can be precisely determined. *Drosophila* has been extensively studied in the fields of genetics and developmental biology. Accurate tracking and recording of its life cycle could significantly promote the understanding of various aspects of biology, agriculture, and medicine.

Noteworthy, no recent improvements have been made in measuring the timing of the developmental transitions in *Drosophila*. Until now, the timing of each developmental stage was manually determined by counting the number of flies at each stage in each vial over 2, 6, 12, or 24 h(*Buhler et al., 2018, Demay et al., 2014, Kulshammer et al., 2015, Nikhil et al., 2016, Yun et al., 2017*); however, this technique has certain limitations. Increasing the temporal resolution is difficult as it would require intensive labor, preventing the detection of subtle changes in transition timing for each event. Moreover, manual counting is neither practicable for large-scale/high-throughput screening nor feasible for identifying multiple phenotypes in individual flies in transition events. In contrast, it does help elucidate the associations between developmental stages and the genetic and environmental factors affecting them. Recent studies reported the use of video cameras to measure developmental timing(*Hironaka et al., 2019*). However, this method is labor-intensive, requires long analytical periods, and is unsuitable for high-throughput analysis.

Here, we present a new scalable method that automatically determines multiple transition time points, such as pupariation, eclosion, and death, of individual fruit flies implementing a basic flatbed CCD scanner. We placed a single fly in each well of a 96- or 384-well microplate, acquired time-lapse images, and analyzed the output with our novel algorithm Sapphire. This system performs automatic developmental analyses at high temporal resolution and could be used in high-throughput gene and chemical screening and analyses of the effects of genetic and environmental factors on development.

## Results

### Design of the Drosophila Individual Activity Monitoring and Detection System (DIAMonDS)

*Drosophila* develops through four developmental stages (embryo, larva, pupa, and adult), subsequently finishing its own life. The static phases (embryo, pupa, and death) alternate with the dynamic phases (larva and adult; Figure 1A). Single-image processing between continuous images distinguishes both phases: dynamic phases are detected as positive signals while static phases simultaneously present no signal (Figure 1B). Therefore, we can precisely identify the transition points between phases by monitoring the static and dynamic status and separately detecting fly phases on all plates captured simultaneously in the same image.

**Figure 1.**
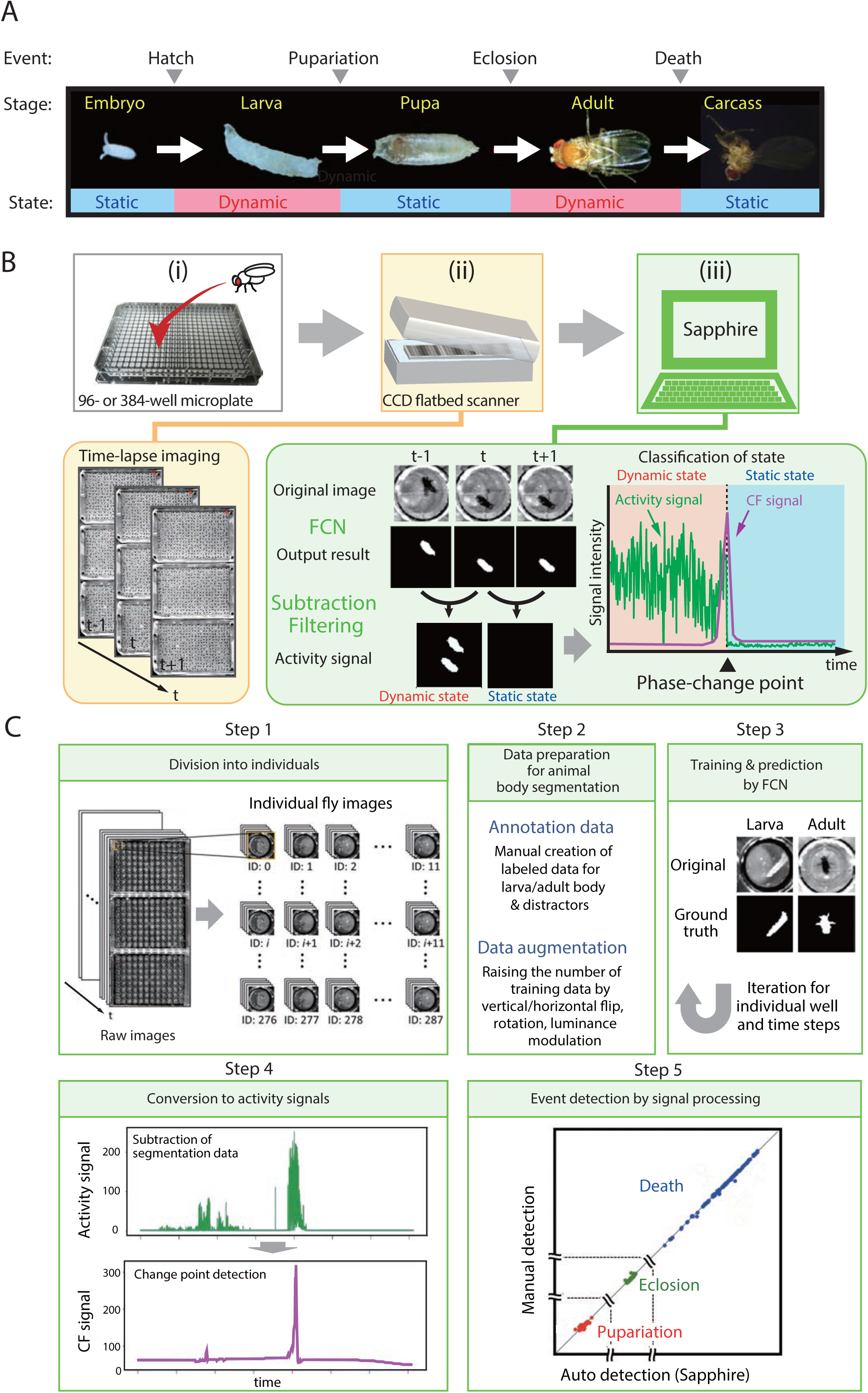
Overview of DIAMonDS. **(A)** Diagram of developmental stages and activity states in *Drosophila melanogaster.* **(B)** Schematic representation of the DIAMonDS procedure. DIAMonDS consists of: (i) microplate preparation; (ii) time-lapse imaging with CCD scanner; and (iii) data analysis by Sapphire. In step (iii), Sapphire calculates activity signal (green line) intensity via animal body detection from FCN images and subtraction processing of every two consecutive images and then determines the CF signal (purple line) from the activity signal. **(C)** Flowchart of Sapphire. Algorithm includes extraction of individual animals from population images (Step 1), training data preparation and augmentation (Step 2), training through data and animal body segmentation (Step 3), segmentation data signaling by subtracting labeled data and transition point detection algorithm (Step 4), event detection, signal processing, and visualization (Step 5).

DIAMonDS consists of an automated time-lapse imaging system and our novel image analysis software named “Sapphire” (Figure 1B). We used a combination of a flatbed CCD scanner and VueScan software (https://www.hamrick.com), which enables multiple scanner units to capture images continuously at certain intervals(*Smith et al., 2014*). To identify the phase change time points for each fly event, single flies are inserted into each well of a 96- or 384-well microplate containing suitable fly media. Up to three microplates are then set on the scanner surface, and time-lapse images are acquired at appropriate intervals until the fly event is completed. A single scanner can monitor 288 or 1,152 individuals in three 96- or 384-well microplates (at the time of death only), respectively. Sapphire then automatically analyzes and detects the transition points of pupariation, eclosion, and death in the newly acquired time-lapse images (Figure 1B,C). Our system can simultaneously monitor multiple scanners with a single personal computer.

As positional bias and fluctuations in environmental factors such as ambient temperature and relative humidity (RH) might markedly reduce the reliability of our system, all experiments were conducted in a plant growth chamber (LPH-410NS) with automatic temperature and humidity regulation and additional USB fans (Supplementary file 6). The scan surface temperature was continuously recorded with button-sized temperature data loggers (NK Labs LLC, Cambridge, MA, USA). Temperatures widely varied among locations under alternating light-dark conditions possibly because of irregular and uneven irradiation. Thus, we acquired time-lapse images under constant darkness to maintain a steady temperature (Figure 1-figure supplement 1).

### DIAMonDS software: Sapphire

Sapphire automatically determines the static-to-dynamic and dynamic-to-static phase changes for all flies according to the time-lapse images acquired by the aforementioned scanner system. The following four processes were implemented in Sapphire to enable automatic life event detection based on individual *Drosophila* images (Figure 1C):

1. Images of single animals were segregated by image processing (Figure 1C, Step 1), enabling the system to readily target individual flies.
2. Semantic segmentation was performed to capture each insect in each well (Figure 1C, Step 2). Recently, artificial intelligence techniques have substantially improved both image classification(*Goodfellow et al., 2016*) and segmentation(*Badrinarayanan et al., 2017, Ronneberger et al., 2015*). Here, we designed a fully convolutional network (FCN) specifying larval and adult segmentation. The FCN has encoder-decoder architecture (Figure 1-figure supplement 2) comprising three blocks, each including a convolution layer with a 3 × 3 filter, up/down sampling layer with a 2 × 2 filter, and dropout layer with a 25% ratio. The encoder and decoder parts were mutually connected by two convolution layers with 3 × 3 filters. The convolution layers in the encoder and decoder were fitted with rectifier linear units. The inputs to the layers were applied by batch normalization. The output layer was also a convolution layer and included reshaping and softmax functions. The annotation data were manually created for larval and adult *Drosophila* and increased by general data augmentation techniques such as vertical and horizontal flips, rotation, and luminance modulation (Materials and methods). After training with augmented data, the segmentation inference was calculated as a probability. If it was > 0.5, the system regarded it as the target region (animal body). If it was < 0.5, the system treated it as background. Consequently, the system obtained binary images wherein the animal body corresponded to 1 and the background was described as 0. Semantic segmentation was applied to all individuals and every sequential population image. All trainings and inferences were performed on a Linux PC (Ubuntu) with four GPUs (GTX 1080Ti). All scripts were written in Python and deep learning libraries such as keras (v. 2.0.9) and tensorflow-gpu (v. 1.4.0).
3. The system converted the labeled data to time series data by subtracting every two consecutive labeled images (Figure 1C, Step 3). After signaling, the system applied the ChangeFinder (CF) algorithm developed to detect turning points(*Takeuchi and Yamanishi, 2006*) and evaluated total individual animal activity from the central gravity distance of the segmented area between two consecutive images (Supplementary Video 1).
4. The system automatically determined life event transitions with single-animal resolutions from the CF signals (Figure 1C, Step 4). Life event transitions were estimated based on the transition points from dynamic to static or vice-versa. These were designated the maximum points in the CF signals. Death and pupariation corresponded to maximum CF points reflecting dynamic-to-static phase changes. In contrast, eclosion was characterized as a maximum CF point corresponding to a static-to-dynamic change. After transition point determination, the system summarized the automatic detection results using various visualization styles (Figure 1C, Step 5 and Figure 1-figure supplement 3). All algorithms and visualization tools were user-friendly web applications developed with the Dash framework in Python (Figure 1-figure supplement 3).

### Pupariation and eclosion timing detection by DIAMonDS

Apparent dynamic-to-static and vice-versa phase changes occur at pupariation and eclosion (Figure 2A). We attempted to detect individual fly pupariation and eclosion with DIAMonDS to validate it. Newly hatched first-instar (L1) larvae were placed into 96-well microplates containing 100 µL well^-1^ standard fly media (Figure 2-figure supplement 1). Three microplates were fixed to the scanner surface, and the scanner was placed inverted in the incubator to prevent fly media leakage. The scanner was powered on, and time-lapse scanning was performed with VueScan (Materials and methods) until all flies eclosed. Thence, time-lapse images were analyzed with Sapphire (Figure 2A,B).

**Figure 2.**
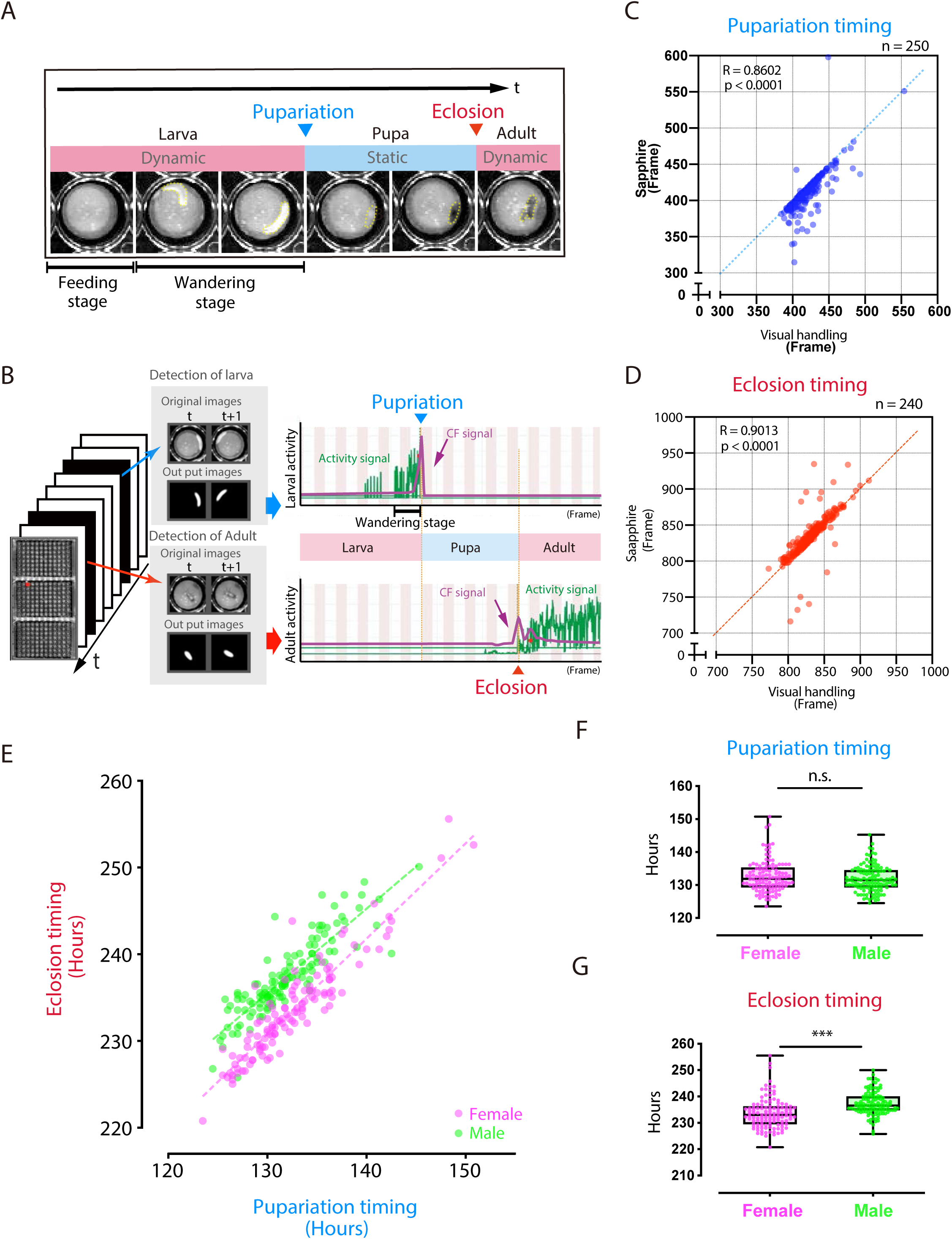
Evaluation of DIAMonDS for automatic detection of individual fly pupariation and eclosion. **(A)** Schematic representation of pupariation and eclosion. Drastic changes in dynamic-to-static and static-to-dynamic states occur at pupariation and eclosion, respectively. At late L3, larval activity increases and transitions from feeding to wandering behavior. Dotted circles indicate animal bodies. **(B)** Time-lapse imaging was conducted until all flies eclosed into adults. Individual pupariation and eclosion transition points were separately analyzed in Sapphire. Wandering L3 larva showing high activity immediately before pupariation. **(C,D)** Scatterplot analyses comparing data for pupariation (**C**) and eclosion (**D**) obtained by Sapphire and visual handling to validate accuracy. **(E-G)**, Scatterplot between pupariation and eclosion timing of individual flies (**E**) and box plots of pupariation (**F**) and eclosion (**G**) timing in males (n = 122; green dots) and females (n = 118; magenta dots). Whiskers indicate minima and maxima (\*\*\**p* < 0.001; n.s., no significant difference; unpaired *t-*test).

The L1 larvae dove into the media until mid-L3 (∼2 d). As time-lapse scanning could not detect larval movements here, this so-called feeding stage was designated as static. In contrast, the larvae typically left the media and moved around the well surface during late L3 (Figure 2A). This so-called wandering stage was designated as dynamic, as there were consecutive high-activity wandering larval signals (Figure 2B), and its duration was 12–24 h. Thereafter, L3 larval activity gradually decreased, and pupariation followed. During the pupal stage (∼100 h), the activity was not detected in a static phase. The second signal wave activity was detected just after dynamic eclosion.

Sapphire independently and automatically detects maximum peaks at the pupariation and eclosion transition points using machine learning based on an FCN to detect larval and adult animals separately (Figures 1C and 2B). To verify Sapphire accuracy, we compared its data against those that were manually derived (visual handling) from the same images. The Sapphire and manual data were nearly identical (Figure 2C,D), with a slight difference of ±10 frames between them for 73.8% and 84.2% of the Sapphire values (Figure 2-figure supplement 2). Film surface contamination after long-term rearing in small wells might have accounted for the observed decrease in Sapphire detection accuracy. We transferred black pupae to new microplates, resumed time-lapse scanning, and detected the eclosion time points. The results of the latter assay clearly demonstrated that Sapphire accuracy was greatly improved relative to that for the experimental setup in which the same microplate was used both for pupariation and eclosion (Figure 2-figure supplement 3).

Sapphire analyzes consecutive images by both fully automated (CF) and semi-automatic (TH) methods (Materials and methods). We compared CF and TH using the same time-lapse image set (Figure 2-figure supplement 3). The output of CF was superior to that of TH, as the former is relatively more sensitive to phase shifts. Thus, Sapphire is functional and highly invaluable in automatic analyses.

The relationship between pupariation and eclosion determined by DIAMonDS clearly revealed sexual dimorphism during development (Figure 2E). On average, females eclosed 4 h earlier than males. A previous study corroborated this observation(*Bainbridge and Bownes, 1981*). The pupariation transition time points did not differ between sexes. Therefore, sexual dimorphism in the eclosion time points reflects the difference in pupal duration between sexes (Figure 2E-G). No significant differences were found among the three 96-well microplates used here, indicating a minimal scan surface positional effect in DIAMonDS during the detection of the pupariation and eclosion time points (Figure 2-figure supplement 4). Altogether, these results indicate that DIAMonDS is suitable for automatic measurement of both the pupariation and eclosion time points at high temporal resolution.

### DIAMonDS enables autonomic measurement of pupariation and eclosion timing at high temporal resolution for each individual

DIAMonDS showed excellent performance in large-scale pupariation timing analyses (Figure 3-figure supplement 1). To evaluate its performance, we explored whether two distinct genetic and environmental conditions affect larval development. First, we used larvae with delayed pupariation at 29 °C (genotype: *R29H01-GAL4*>*UAS-TeTxLC*), in which *tetanus toxin light chain* (*TeTxLC*) impaired serotonergic SE0_PG_ neuron activity(*Shimada-Niwa and Niwa, 2014*). To evaluate the influences of initial larval age on DIAMonDS measurement accuracy, we used L2, early L3, and late L3 larvae in the microplate assays. Using only a few specimens, DIAMonDS successfully detected pupariation delays in larvae of all ages (Figure 3A). Thus, DIAMonDS performance remains highly stable over a wide range of conditions.

**Figure 3.**
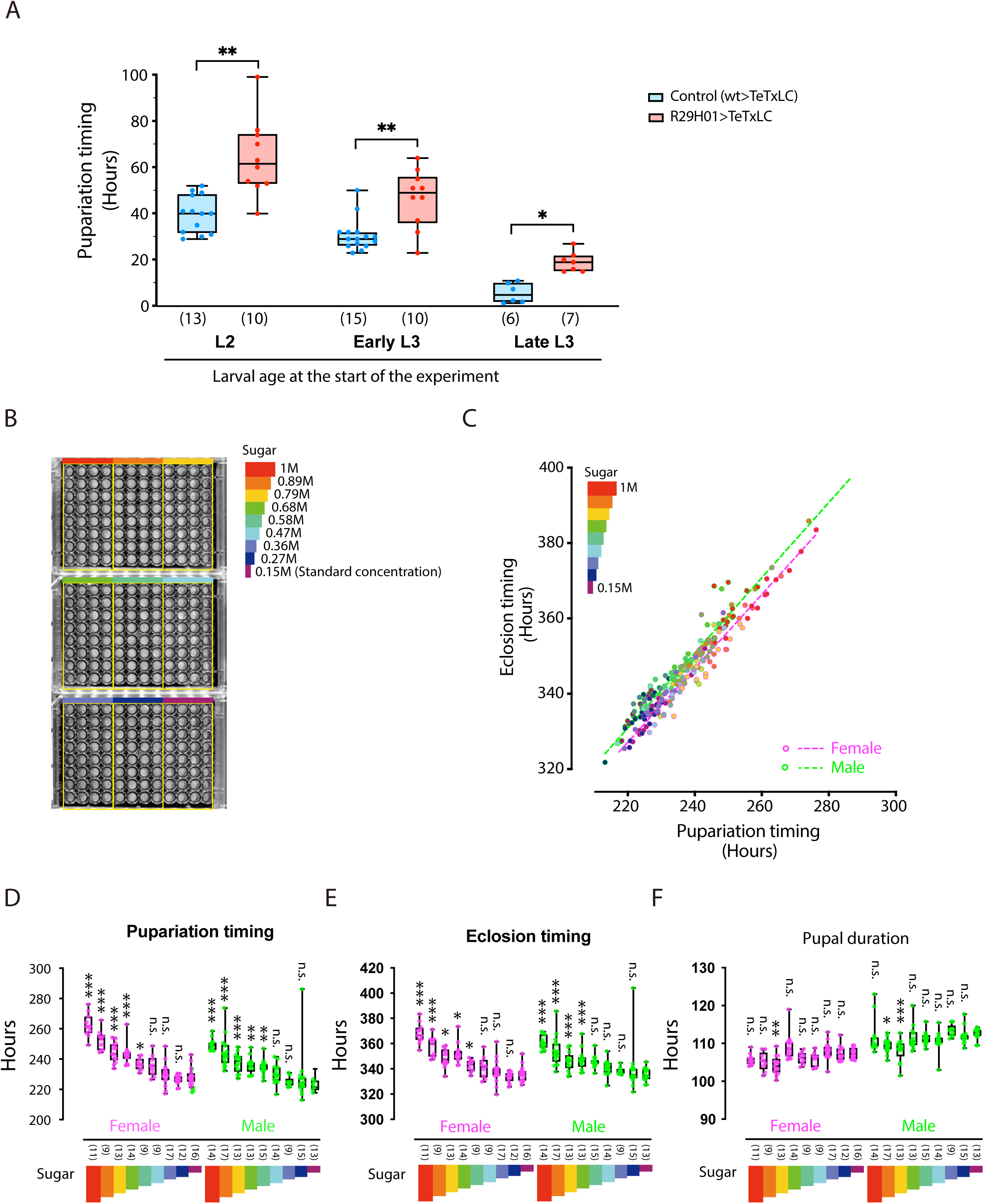
DIAMonDS is suitable for phenotypic analyses during larval development. **(A)** Box plot analysis of pupariation timing in flies with impaired ecdysteroid biosynthesis at 29 °C (genotype: *R29H01>TeTxLC; R29H01>+* as a control). Larval age (L2, early L3, and late L3) at start of measurement had negligible impact on DIAMonDS analysis accuracy. Y-axis indicates pupariation timing from start of experiment. Number of flies analyzed indicated in parentheses on the graph. Whiskers indicate minima and maxima (\**p* < 0.05; \*\**p* < 0.01; \*\*\**p* < 0.001; n.s., no significant difference; multiple *t*-test). **(B)** Three 96-well microplates were subdivided into nine regions according to sucrose concentration (0.15–1 M) in media used for DIAMonDS. **(C)** Scatterplot between pupariation and eclosion timing in males and females. **(D-F)** Box plots of pupariation timing (**D**), eclosion timing (**E**), and pupal duration (**F**). Whiskers indicate minima and maxima (\**p* < 0.05; \*\**p* < 0.01; \*\*\**p* < 0.001; n.s., no significant difference vs. standard diet group; one-way ANOVA followed by Dunnett’s multiple comparison test). Number of flies analyzed indicated in parentheses on the graph.

We also used DIAMonDS to clarify the relationship between sugar concentration and development in the *w^1118^* fruit fly strain (Figure 3B-F). A previous report stated that *Drosophila* larvae that were administered a high-sugar diet presented a type 2 diabetes-like phenotype and developmental delay(*Musselman et al., 2011*). L1 larvae were placed in wells containing normal to high sugar concentrations, scanned by time-lapse imaging, and the pupariation and eclosion time points were detected by Sapphire. Both pupariation and eclosion were gradually delayed in a sugar concentration-dependent manner. Therefore, excess sugar adversely affects larval growth (Figure 3B-E). When we compared the pupariation and eclosion time points among individual larvae, the pupal durations were nearly constant regardless of sugar concentration. Thus, determination of the pupal period is dependent on larval sugar diet (Figure 3F). Interestingly, Northrop reported that prolongation of the pre-imago stage of *D. melanogaster* by yeast supplementation had no impact on pupal stage duration(*Northrop, 1917a, b*). The process by which pupal duration is determined may be independent of larval dietary intake. Overall, DIAMonDS is a powerful toolkit for detecting the pupariation and eclosion time points and discloses the effects of several endogenous and exogenous factors on individual fly development.

### Detection of individual adult death events by DIAMonDS

Adult death time points can be measured at high temporal resolution with DIAMonDS. This tool efficiently detects sudden shifts from a dynamic to a static phase. Depending on the objective, experiments are performed over a broad range of time scales extending to nearly 3 months. Small- and large-scale impact assessment targets may include mutant phenotype, environmental stress, and chemicals and drugs(*Afschar et al., 2016, Harshman et al., 1999, Lin et al., 1998, Parkes et al., 1998, Piper and Partridge, 2016, Tsuda et al., 2010, Ziehm et al., 2015*). DIAMonDS may be implemented using 96- and 384-well microplates to accommodate various parameters in death timing detection. The 96-well type is used in long-term detection, whereas the 384-well plate is better suited for short-term (≤10 d) detection and high-throughput assays.

We collected time-lapse scanning images at 1-min intervals for adult flies (4 d post-eclosion) maintained under starvation conditions (70 µL 1% agar/water (w/v)/well) in order to optimize DIAMonDS for 384-well microplates (Figure 4A). After the image data series consisted of dying individual flies, death time points were determined manually (visual manipulation) and by Sapphire. Both procedures generated nearly identical survival curves and were highly correlated (R = 0.9925), indicating the reliability of Sapphire (Figure 4A–D and Figure 4-figure supplement 1). Next, we examined whether the well positions within a microplate influence the death time points. No significant differences were found between each plate or among the sub-areas within each plate (Figure 4E and Figure 4-figure supplement 2). We also evaluated adult *w^1118^* fly starvation tolerance in 96-well microplates and obtained results similar to those acquired from 384-well microplates (Figure 4F). Therefore, DIAMonDS is highly effective at detecting fruit fly death events.

**Figure 4.**
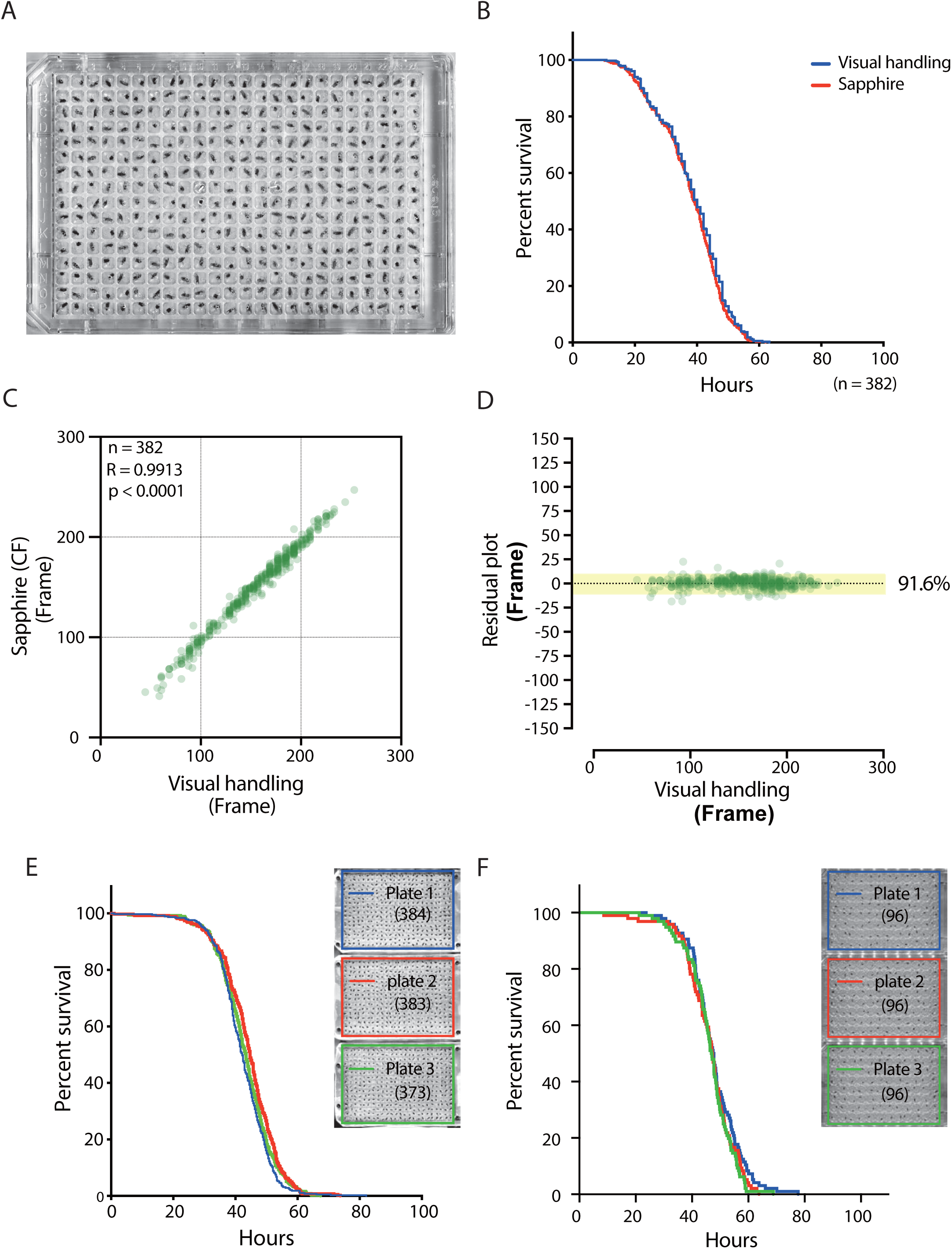
Evaluation of DIAMonDS for detection of death of individual adults. **(A)** A 384-well microplate with flies prepared for DIAMonDS. **(B)** Survivorship curves plotted by Sapphire (red line) or visual handling (blue line) using the same data. **(C,D)** Scatterplot and residual plot analysis comparing Sapphire (CF method) and visual handling to validate accuracy (n = 382). **(E,F)** Survivorship curves for starvation resistance tests on adult male *w^1118^* flies using three 384- (**E**) and three 96-well (**F**) microplates. Number of flies analyzed indicated in parentheses in each plate.

### DIAMonDS performs stress resistance assays with high temporal resolution

To assess the efficacy of DIAMonDS at evaluating fly survival under various stress conditions, we performed a starvation assay on male and female *w^1118^* flies (Figure 5A). They presented sexual dimorphism in terms of starvation resistance. Moreover, the temporal resolution used here (15 min intervals) was higher than those reported in previous studies (Figure 5B and Figure 5-figure supplement 1A,B)(*Gronke et al., 2010, Li et al., 2018*).

**Figure 5.**
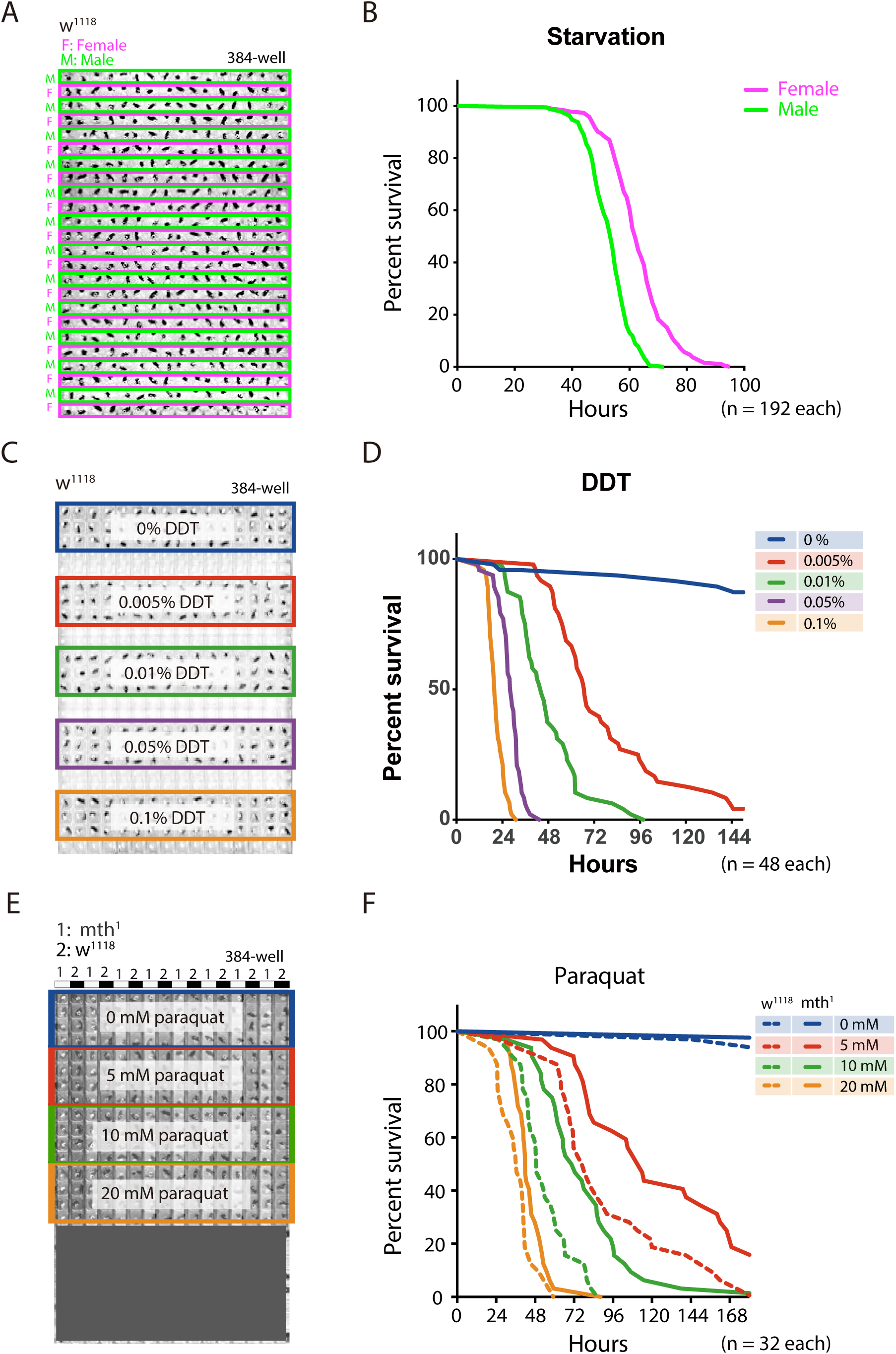
Adult survival curve detection by DIAMonDS. **(A,B)** Starvation tolerance test using male and female *w^1118^* flies (n = 192 each). Rows of males and females were alternately arranged in 384-well microplate (**A**). Representative male and female survivorship curves (**B**). **(C,D)** DDT resistance test on male *w^1118^* flies. A 384-well microplate with YS media containing DDT concentration series (0–0.1%). Forty-eight male flies were exposed to each concentration and subjected to DIAMonDS (**C**). Survivorship curves show concentration-dependent toxic effects of DDT (**D**). **(E,F)** Paraquat resistance test on male *mth^1^* and *w^1118^* flies (n = 32 each). A 384-well microplate containing media with paraquat (0, 5, 10, and 20 mM). Rows of *mth^1^* and *w^1118^* flies were alternately arranged in wells (**E**). Survivorship curves for *mth^1^* mutants (solid lines) and *w^1118^* flies (dotted lines) substantially differed at all paraquat concentrations (**F**).

We then subjected the flies to various concentrations of dichlorodiphenyltrichloroethane (DDT) to identify possible resistance (Figure 5C,D)(*Afschar et al., 2016*). To a 384-well microplate, we added media consisting of 5% (w/v) sucrose and 0.5% (w/v) yeast as described in Supplementary file 1. The 48 *w^1118^* males in each well were exposed to 0, 0.005, 0.01, 0.05, or 0.1% DDT (Figure 5C). DIAMonDS showed distinct variance in DDT resistance even between only slightly differing DDT concentrations (Figure 5D and Figure 5-figure supplement 1C,D).

We also investigated whether DIAMonDS can detect mutant fly phenotypes in stress resistance assays. Flies with the loss-of-function mutation *methuselah* (*mth*) present extended lifespan and resistance to several stressors including paraquat(*Lin et al., 1998*). To test whether DIAMonDS can discriminate *mth^1^* mutant stress tolerance phenotypes, we backcrossed the *mth^1^* mutant five times with *w^1118^* and compared the responses of both types of males to different paraquat concentrations (Figure 5E,F). The *mth^1^* mutant presented significantly greater resistance to several paraquat concentrations than *w^1118^*. The results of this assay demonstrated that paraquat resistance could be detected in the *mth^1^* mutant using lower paraquat concentrations than those tested in an earlier study (20 mM)(*Lin et al., 1998*). Thus, DIAMonDS can readily identify optimal concentrations at high temporal resolution in drug resistance assays.

### Sequential detection of multiple life events (pupariation, eclosion, and lifespan) using DIAMonDS

DIAMonDS successfully measured lifespans for individual *w^1118^* flies. For long-term analysis, flies must be transferred to new microplates. We performed sequential pupariation, eclosion, and lifespan measurements for individual *w^1118^* as shown in Figure 6A-C. The mean lifespans calculated and survival curves plotted for individual males and females were consistent with those of previous reports (Figure 6C)(*Liu et al., 2009, Schriner et al., 2014, Suh et al., 2008, Trostnikov et al., 2019*).

**Figure 6.**
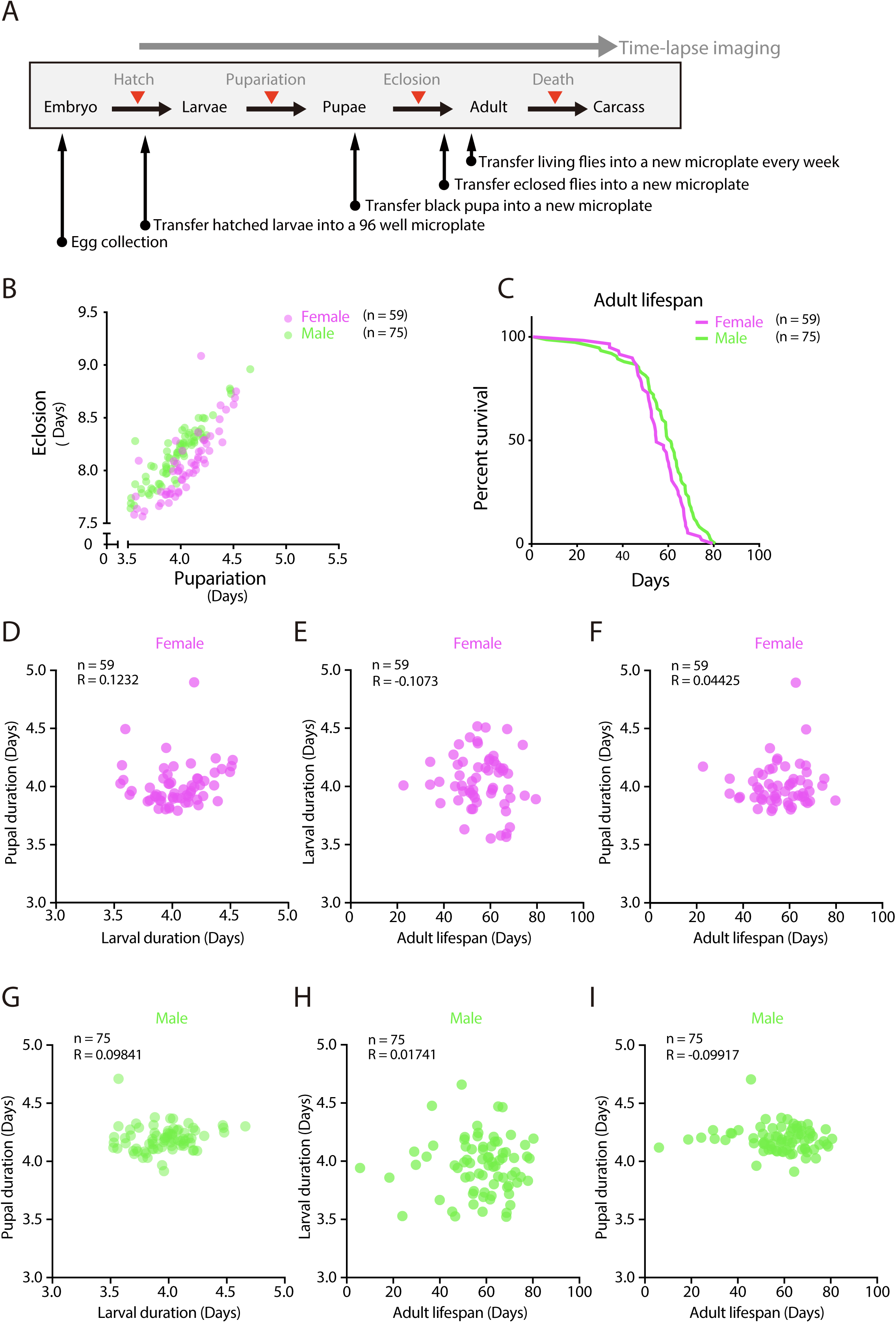
Tracing entire life events of individual flies by DIAMonDS. **(A)** Schematic diagram showing detection of pupariation, eclosion, and death transition points for each individual in DIAMonDS. **(B)** Scatterplot between pupariation and eclosion timing for individual female (n = 59) and male (n = 75) flies. **(C)** Survivorship curves for adult female (n = 59) and male (n = 75) flies. **(D-I)** Pearson’s correlation scatter plots indicate relationships between larval and pupal duration (**D**,**G**), between larval duration and adult lifespan (**E,H**), and between pupal duration and adult lifespan (**F,I**). Data are separately presented for female (**D-F**, n = 59 each) and male (**G-I**, n = 75 each) flies.

Several studies described the relationships between developmental time and adult lifespan among various species(*Marchionni et al., 2020*). Certain reports showed that prolonging developmental duration by yeast supplementation did not affect lifespan(*Hunter, 1959, Northrop, 1917b*). However, few studies focused on the associations between developmental timing and adult lifespan in *Drosophila*. To establish whether developmental stages and adult lifespans are interdependent in individual flies, we analyzed the following relationships: larval and pupal duration (Figure 6D,G), larval duration and adult lifespan (Figure 6E,H), and pupal duration and adult lifespan (Figure 6F,I). No significant correlations between the data for these pairs of parameters in either sex (Figure 6D-I) and those reported in previous studies were found. As the flies had the same genetic background and were exposed to the same diet and environmental conditions throughout all their life stages, it was unlikely that their genotypes influenced the relationship between development time and lifespan, demonstrating that DIAMonDS can detect unique relationships between developmental time and lifespan under different intrinsic and extrinsic conditions.

In this study, we were unsuccessful at using DIAMonDS to run long-term lifespan assays on other wild type fruit fly strains such as *Oregon-R* and *Canton-S*. Lethality accidentally escalated because the rearing conditions deteriorated. For instance, water droplets condensed and coalesced on the plate surfaces. However, these errors might have reflected relative metabolic and/or genetic differences among fly strains. Optimization of the experimental conditions will be necessary to effectively use DIAMonDS to conduct lifespan tests on a wide range of fly strains.

## Discussion

In this study, we attempted to develop DIAMonDS as a new tool to analyze automatically and sequentially the transition time point in the growth and developmental multi-stage of individual flies and use the transition time points as a phenotype. We demonstrated that DIAMonDS determines *Drosophila* pupariation, eclosion, and death at high temporal resolution. Further, it can analyze the relationships among stages by sequentially detecting multiple life events in many individuals. Thus, DIAMonDS can help clarify the complex interactions among genetic and environmental factors throughout the *Drosophila* life cycle. DIAMonDS can also eliminate the constraint of long data acquisition and analysis time intervals, operate multiple scanners simultaneously, and facilitate high-throughput analysis. Overall, DIAMonDS substantially ameliorates *Drosophila* research endeavors compared to conventional manual counting methods.

DIAMonDS automatically detects time points from time-lapse images via the novel algorithm Sapphire. Our results indicated that relative to manual detection, DIAMonDS correctly detected 74–85% of the pupariation and eclosion and ∼92% of the death events within ±10 frames. Thus, DIAMonDS is fully functional both for preliminary experiments and large-scale screening. The output of Sapphire can guide the manual determination of exact values. Therefore, DIAMonDS can generate publishable high-quality data. Detection accuracy could be greatly improved by reducing noise in time-lapse image acquisition. Data quality can also be enhanced via the machine learning training step in Sapphire. Optimization of experimental conditions is an important initial prerequisite step for stable, highly reliable data acquisition.

Nevertheless, DIAMonDS has certain limitations. First, it is unsuitable for the analysis of insects with normal circadian rhythms as the flatbed scanners repeatedly emit light for imaging. Second, the confined well space used for insect rearing might adversely affect individual fly health and behavior. Third, this approach might be inappropriate for establishing the effects of volatile or unstable substances on flies. Fourth, the reliability of this system may be reduced as the surfaces of the microplate lids become dirty over time. These problems could be solved by further improvement in the system. For example, detection of the circadian rhythm might be achieved by image acquisition using infrared rays, and It seems that the use of large-wells might expect to reduce the adverse effects of rearing on small-wells.

To date, we successfully used DIAMonDS to analyze the normal lifespan of the *w^1118^* fruit fly strain but not those of the wild type *Oregon-R* and *Canton-S.* One reason for this constraint is the frequent occurrence of accidental death caused mainly by water droplet coalescence on the microplate wall surfaces. Differences between strains may be due to differences in genetic background or intestinal environment. The experimental setup will require further optimization to overcome these limitations in lifespan analysis by DIAMonDS.

Here, we reported a novel technique for measuring multiple life events and their transition time points in *Drosophila*. In principle, DIAMonDS can also determine life cycle phase shifts in other small animals, and we intend to expand its applicability in medical, agricultural, and ongoing biological research.

## Materials and methods

### *D. melanogaster* stocks and rearing conditions

The wild type strain used here was mainly *w^1118^*. The *mth^1^* (BDSC #27896) mutant males were used in the paraquat resistance test after backcrossing six-fold with *w^1118^*. *R29H01-GAL4* (BDSC #47343) and *UAS-TeTxLC* (BDSC #28838) were obtained from the Bloomington Drosophila Stock Center, Bloomington, IN, USA. All fly strains were maintained on standard medium at 25 °C (Supplementary file 1).

Most experiments were conducted in a plant growth chamber (LPH-410NS; NK System Co. Ltd., Nagoya, Japan) maintained at 25 °C and 60% RH. For the experiment using *R29H01-GAL4>UAS-TeTxLC,* the larvae were reared at 29 °C to increase GAL4 activity.

### DIAMonDS hardware and layout

Detailed DIAMonDS hardware and layout are described in Supplementary file 6.

### Microplate preparation for DIAMonDS

Microplates with 96 or 384 wells were filled with standard fruit fly culture media (Supplementary file 1). A handmade acrylic lid was used for the 384-well microplate (Supplementary file 2). Titer stick film (Watson, Tokyo, Japan) was used for the 96-well microplate (Supplementary file 3). To determine the pupariation and eclosion timings, the film seal on the 96-well microplate was perforated with air holes (0.35 mm diameter; eight holes/well) over each well. For the stress resistance and lifespan tests, the 96-well microplate was sealed with film, which was then cut to form a cross shape over each well. Individual adult flies were transferred to each well without anesthesia. For the 384-well microplate, the flies were anesthetized with triethylamine and placed individually into each well (Supplementary file 4). The acrylic lid was then securely affixed to the microplate with screws and masking tape (Supplementary file 2).

### Time-lapse image acquisition

Three microplates were secured with glass slides to the flatbed scanner surface as described in Supplementary file 5. The scanner was then connected to a personal computer. “VueScan” scanner software in the PC was launched, and time-lapse images were acquired at 1–15-min intervals (Supplementary file 6).

### Starvation and drug resistance test

For the drug resistance assay, the wells of a microplate were filled with yeast-sucrose medium containing the appropriate chemicals as shown in Supplementary file 1. For the starvation test, the wells of a microplate were filled with 1% (w/v) agar (Supplementary file 1). Virgin adult flies were collected under CO_2_ anesthesia, stored in a vial with normal food for 3–5 d, placed individually into each microplate well, and subjected to time-lapse image acquisition.

### Pupariation and eclosion time point detection for the same individuals

To measure pupariation and eclosion timing, newly hatched L1 larvae (24–25 h AEL) were used as the lethality was too high when embryos were used (Figure 2-figure supplement 1A). The large media volume in each well (150 µL) also increased lethality because the embryos suffocated when the media became very viscous. Therefore, we used 100 µL media for this approach (Figure 2-figure supplement 1B). L2 or L3 larvae were also used in the DIAMonDS experiments. After the larvae were loaded into the microplate wells, the microplate was covered with Titer stick film (Watson, Tokyo, Japan), inverted on a predetermined area of the flatbed scanner surface, and fastened with tape. The scanner was then inverted in the incubator to prevent liquefied culture medium from falling onto the film surface. Time-lapse scanning was then run in VueScan in the PC. Scanning was terminated when all individuals eclosed to adults. The time-lapse images were analyzed in Sapphire (Figure 1B,C).

### Pupariation, eclosion, and lifespan detection in identical individuals in DIAMonDS

Microplates were prepared with synchronized L1 larva (one/well), and images were acquired as previously described. Developmental stages for individuals were analyzed according to the images saved to the PC. After all individuals reached the black pupa stage, the pupae were transferred to the same well positions in a new microplate. The wells were sealed with new film, and scanning was resumed. After all flies completely eclosed, the individual adults were transferred to the same well positions in a new microplate, and scanning was restarted to measure individual lifespans. Adult flies were transferred weekly to new microplates until all flies died. The acquired images were then analyzed in Sapphire. Finally, we introduced the data obtained by Sapphire into the DIAMonDS analysis templates (Supplementary file 9) and calculated the time of each life-event (Supplementary file 7).

### Life event detection algorithm and software: Sapphire

The present system included an automated, high-accuracy life event detection algorithm and image and signal processing (Figure 1C). The software could be obtained from (https://github.com/kanglab/Sapphire/tree/master). The contents in the address also describe the installation, license, and system requirements such as directory tree and external dependencies. Quantitative algorithm summary and robustness in various parameter spaces are described in Supplementary file 8.

### Image processing

The present system performed semantic segmentation via a deep neural network whose architecture is described in Figure1-figure supplement 2. Sufficient annotation data increases inference accuracy in deep learning. For network training, handmade supervised data were prepared and applied to image data augmentation. For adult body segmentation in death detection, the annotation data for 178 adults were introduced and magnified up to 71,200 by general image data augmentation (vertical and horizontal flips, rotation, and luminance modulation). For adult body segmentation in pupariation and eclosion detection, annotation data for 300 adults and 5,760 larvae (as distractors) were prepared and magnified by ≤60,000 and ≤57,600, respectively. For larval body segmentation in pupariation and eclosion detection, annotation data for 1,839 larvae and 183,900 augmented data points were introduced to train the network. The present system detected rough outlines of the animal bodies (Supplementary file 8 and Supplementary file 8A) despite the comparatively small amount of annotation data generated by two datasets.

### Signal processing

In the present system, subtractions between consecutive segmentation images were converted to signals in CF, which is a change point detection algorithm(*Takeuchi and Yamanishi, 2006*).

The CF signal was obtained as follows:

1. Considering the following AR model for time series data *x_t_*:

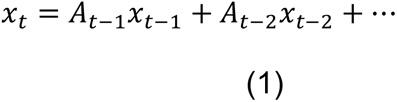
2. Inference of parameters *θ* = (*A*_1_, …, *A_k_*, *μ*, *σ*) by maximizing *I*:

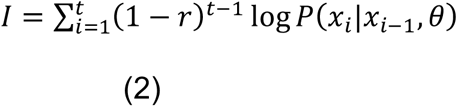
3. Calculation of scores:

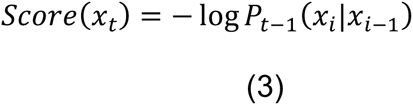
4. Time averaging of scores:

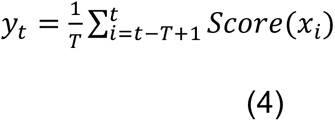
5. Recalculating scores for *y_t_*

For death detection, the CF signal was calculated from the adult body segmentation and the death timing and determined as a maximum CF signal point capturing dynamic-to-static changes (Supplementary file 8 and Supplementary file 8B). On the other hand, two CF signals were obtained from the adult and larval body segmentations in pupariation and eclosion detection. Pupariation is defined as the maximum dynamic-to-static transition point for larvae. Eclosion is defined as the maximum static-to-dynamic transition point for adults (Supplementary file 8 and Supplementary file 8B).

### Statistics

Data were analyzed, and graphs were plotted in GraphPad Prism v. 8 (GraphPad Software, San Diego, CA, USA). A student’s unpaired, two-tailed *t*-test was performed to compare differences between groups in each experiment and Dunnett’s method of one-way ANOVA was used for multiple comparison test (\*\*\**p* < 0.001; \*\**p* < 0.01; \**p* < 0.05; n.s., no significant).

## Acknowledgements

We are deeply grateful to Tadashi Uemura (Kyoto University) for comments, discussion and helpful supports. We thank Yuko Iijima, Eisuke Imura and Hsin Kuang Lin for their technical assistance. We are also grateful to Seino Masaki, Maki Otori, Mayu Kudo, Masaya Hata and Hiroki Watanabe for their technical support in image processing. I received generous support from Yoichi Shinkai (RIKEN CPR) and Shunsuke Ishii (RIKEN CPR). This work was supported by the AMED (Japan Agency for Medical Research and Development) [JP18gm1110001 to K.S., S.K. and R.N.] and JST PRESTO (Japan Science and Technology agency, Precursory Research for Embryonic Science and Technology) [JPMJPR12M5 to K.S.].

## Author contributions

K.S. and S.K. conceived and designed experiments. K.S., S.K. and T.M. designed and implemented hardware and software. K.S. constructed and calibrated equipment. K.S., S.K., T.M., Y.S. and R.N. performed experiments. S.K., T.M., and K.S. designed analytic tools. K.S., S.K. and T.M. analyzed data and interpreted results. K.S. and S.K. wrote the manuscript with contribution from all authors. K.S. and S.K. supervised the project.

## Competing interests

The authors declare no competing financial interests.

## Figure supplement

**Figure 1-figure supplement 1.**
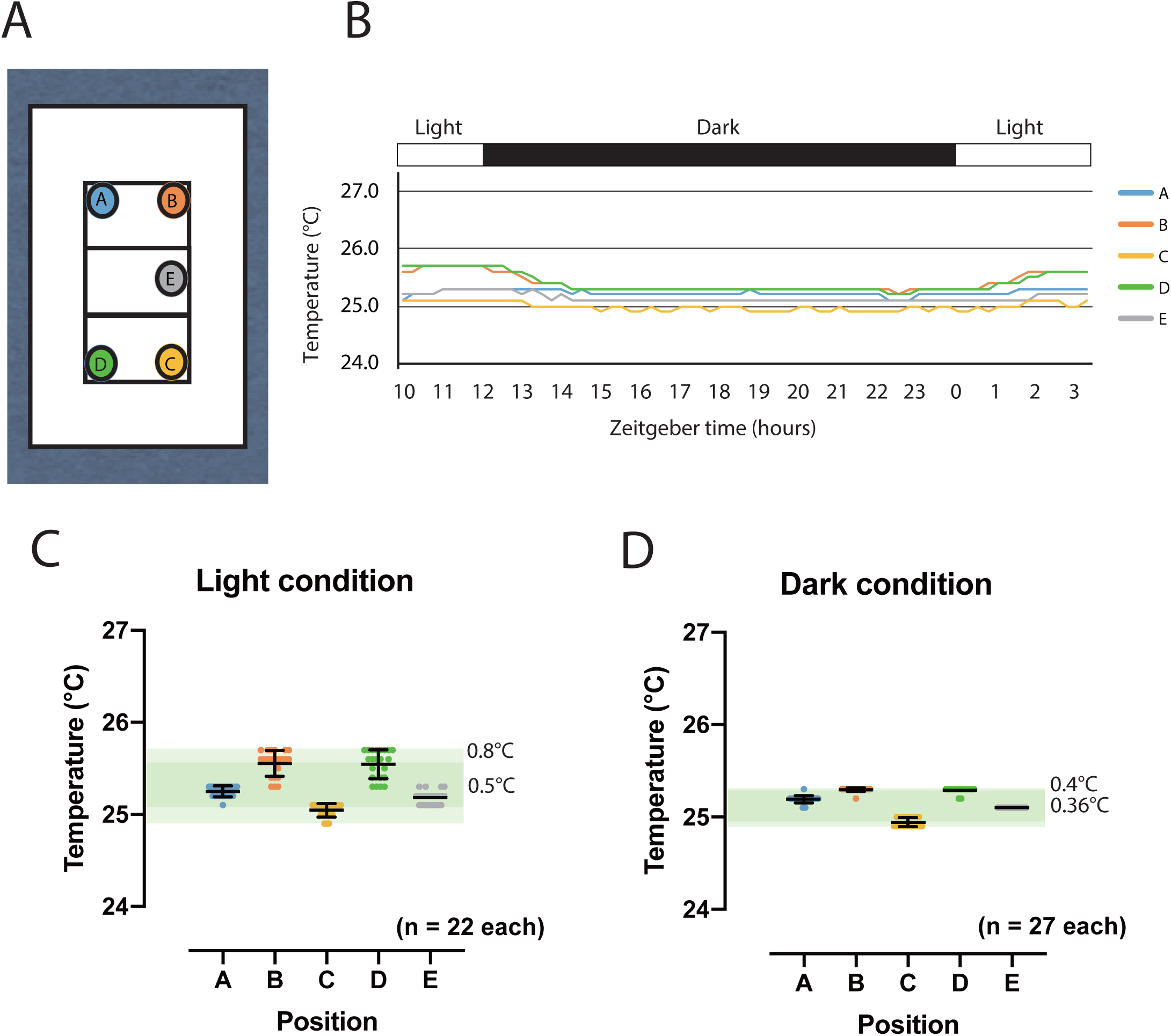
Scanning surface temperature distribution. **(A)** Temperature measurement points on scan surface. Temperature measured by temperature data logger (Thermochron SL type, NK Labs, Cambridge, MA, USA) in high-resolution mode. **(B)** Daily temperature time course under 12:12 light-dark cycle. **(C,D)** Temperature at each time point under light (**C**) and dark (**D**) conditions. Dark and light green bands represent mean and maximum/minimum temperature ranges, respectively.

**Figure 1-figure supplement 2.**
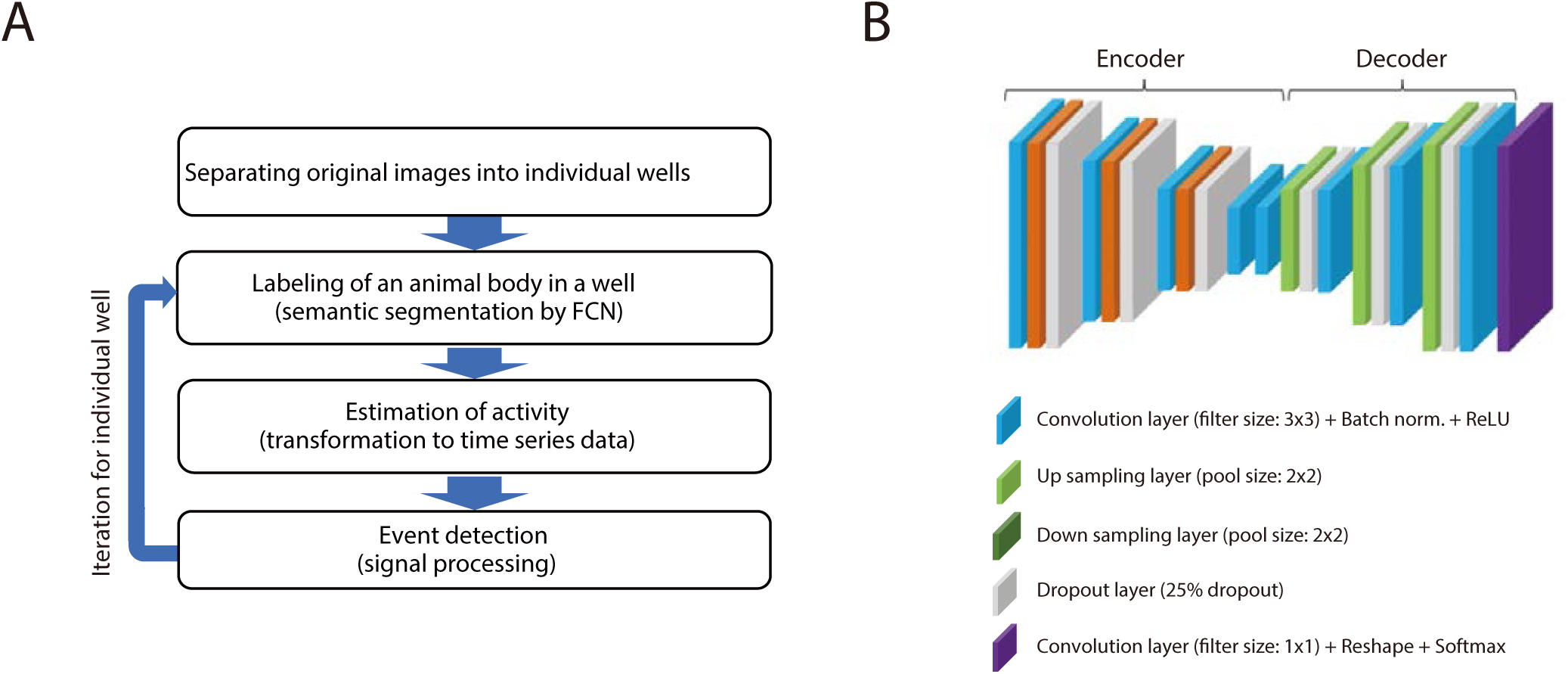
Schematic representation of process of Sapphire and architecture of FCN. **(A)** Automatic life event detection algorithm. Detailed procedure described in Figure 1C. **(B)** Schematic architecture of fully convolutional network (FCN) designed for animal body segmentation.

**Figure 1-figure supplement 3.**
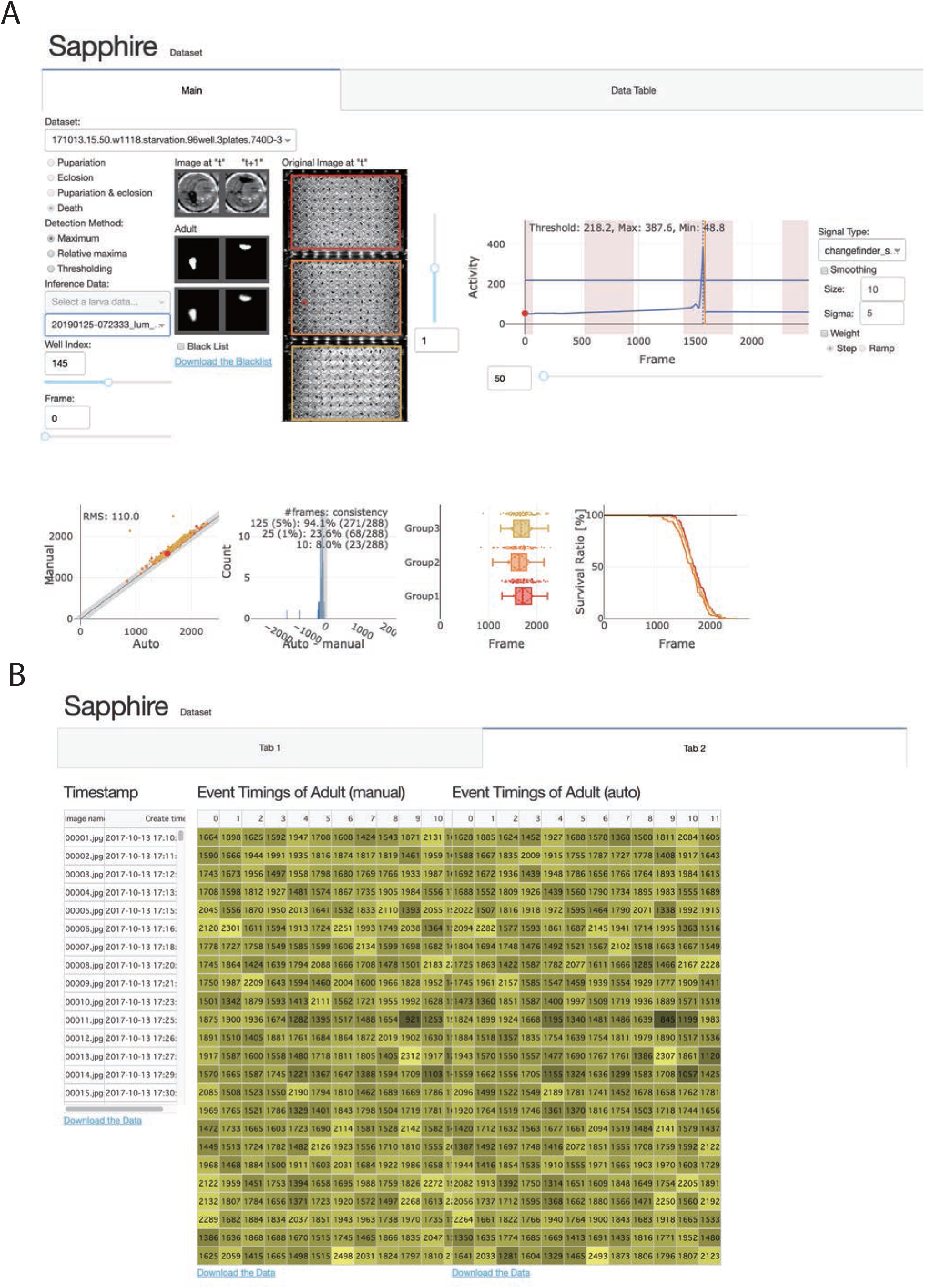
Sapphire user interface. **(A)** Under the tab “Table 1”, appropriate datasets, detection methods, and inference data may be selected. We can also confirm and select well images and segmented areas of two consecutive images. We may confirm the result of Sapphire analysis on the selected well (upper left graph) and change “signal type,” “smoothing,” and “weight.” We can compare data analyzed by Sapphire with those detected manually (lower panels). **(B)** In tab “Table 2”, we can download the timestamp and analyzed data.

**Figure 2-figure supplement 1.**
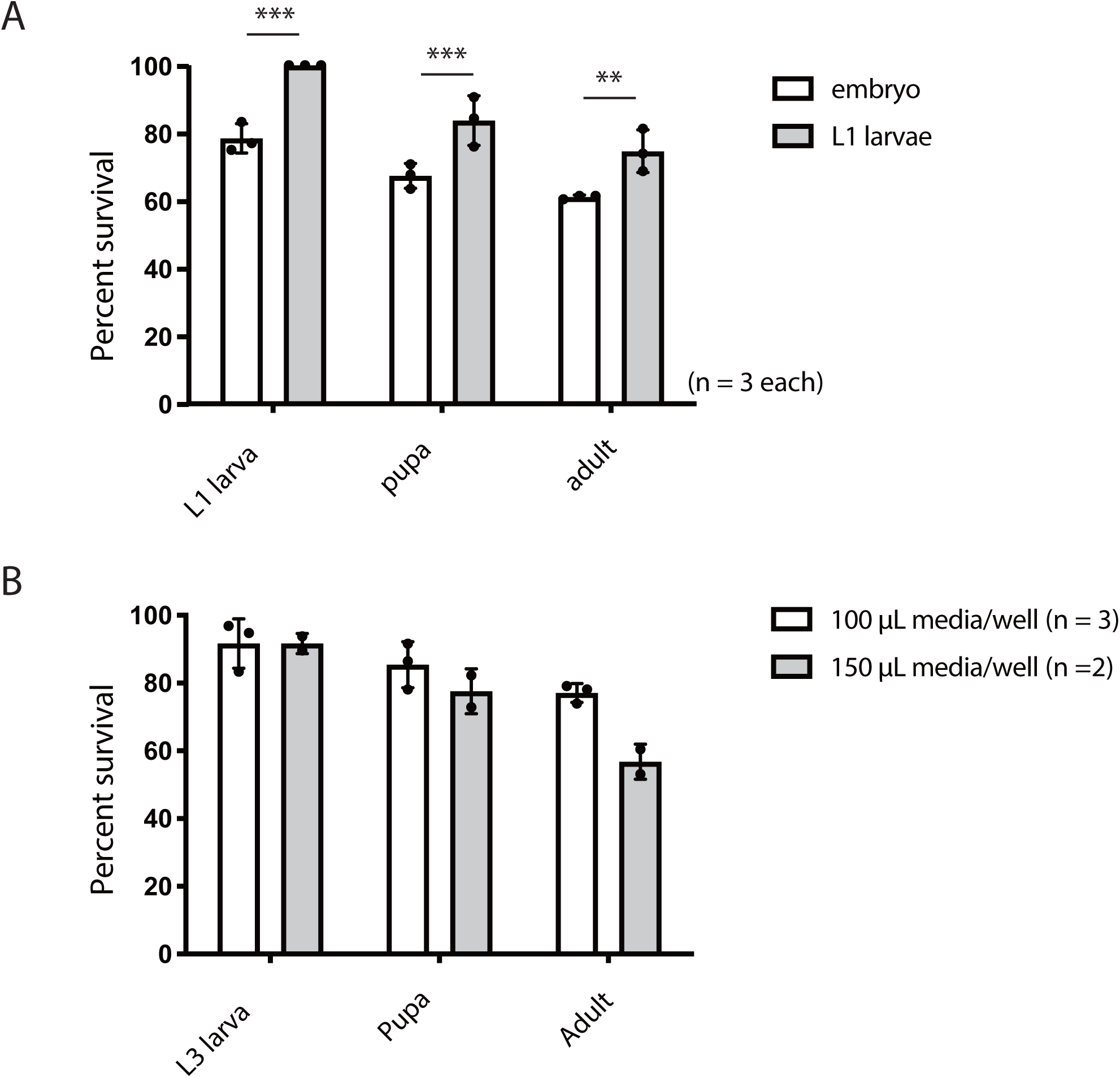
Effects of experimental conditions. **(A)** Percent survival at each developmental stage (1st instar (L1) larva, pupa, and adult stages) when a single embryo or L1 larva is installed in each well of a 96-well microplate. Averages with s.d. are shown (n = 3 each; \*\*\**p* < 0.001; \*\**p* < 0.01; n.s., no significant difference; Student’s unpaired *t*-test). **(B)** Percent survival at each developmental stage (3rd instar (L3) larva, pupa, and adult stages) in 100 or 150 μL medium/well. Averages with s.d. are shown (n = 3 and 2, respectively).

**Figure 2-figure supplement 2.**
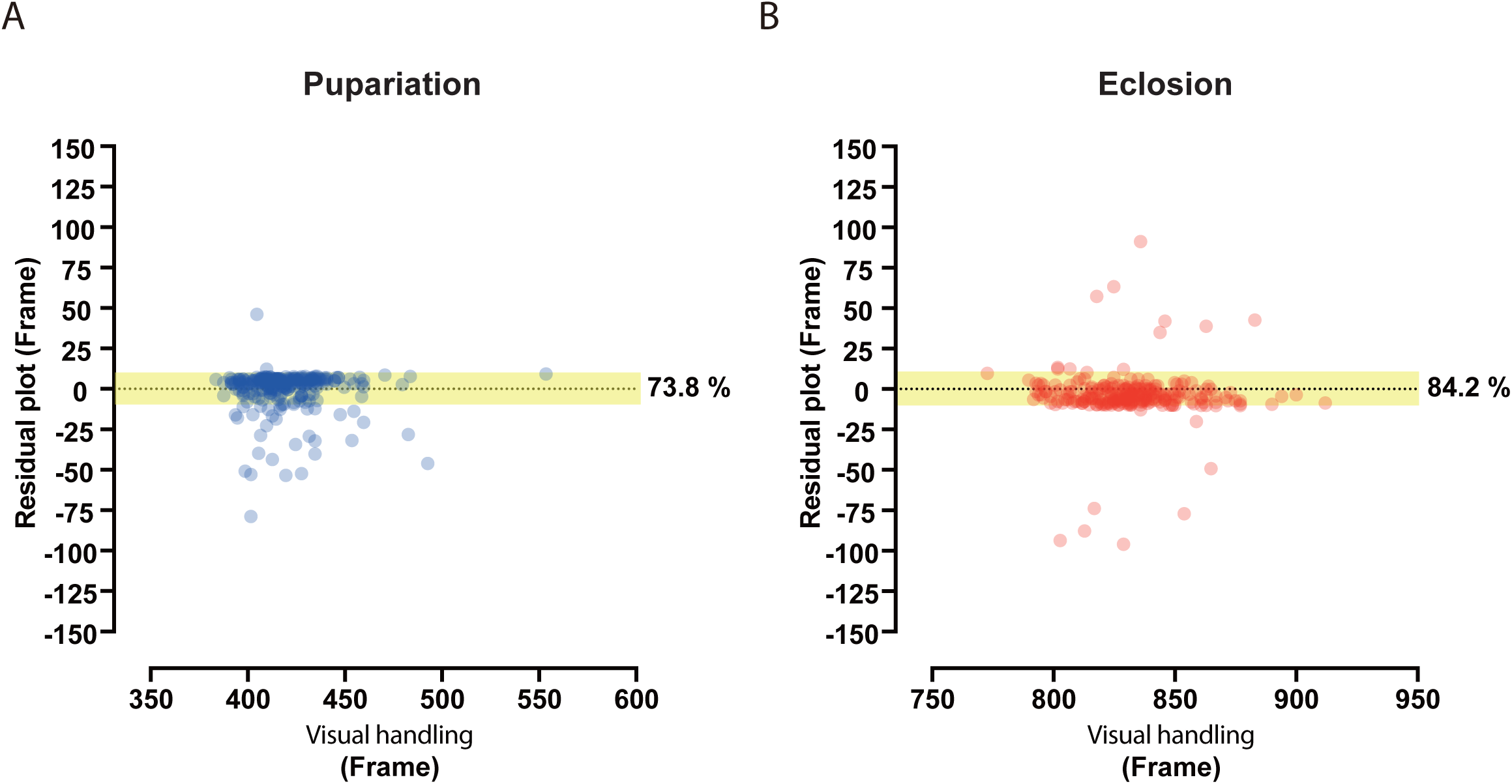
Residual plot analysis of pupariation and eclosion time points. **(A)** Residual plots of pupariation corresponding to Figure 2C. **(B)** Residual plots of eclosion corresponding to Figure 2D.

**Figure 2-figure supplement 3.**
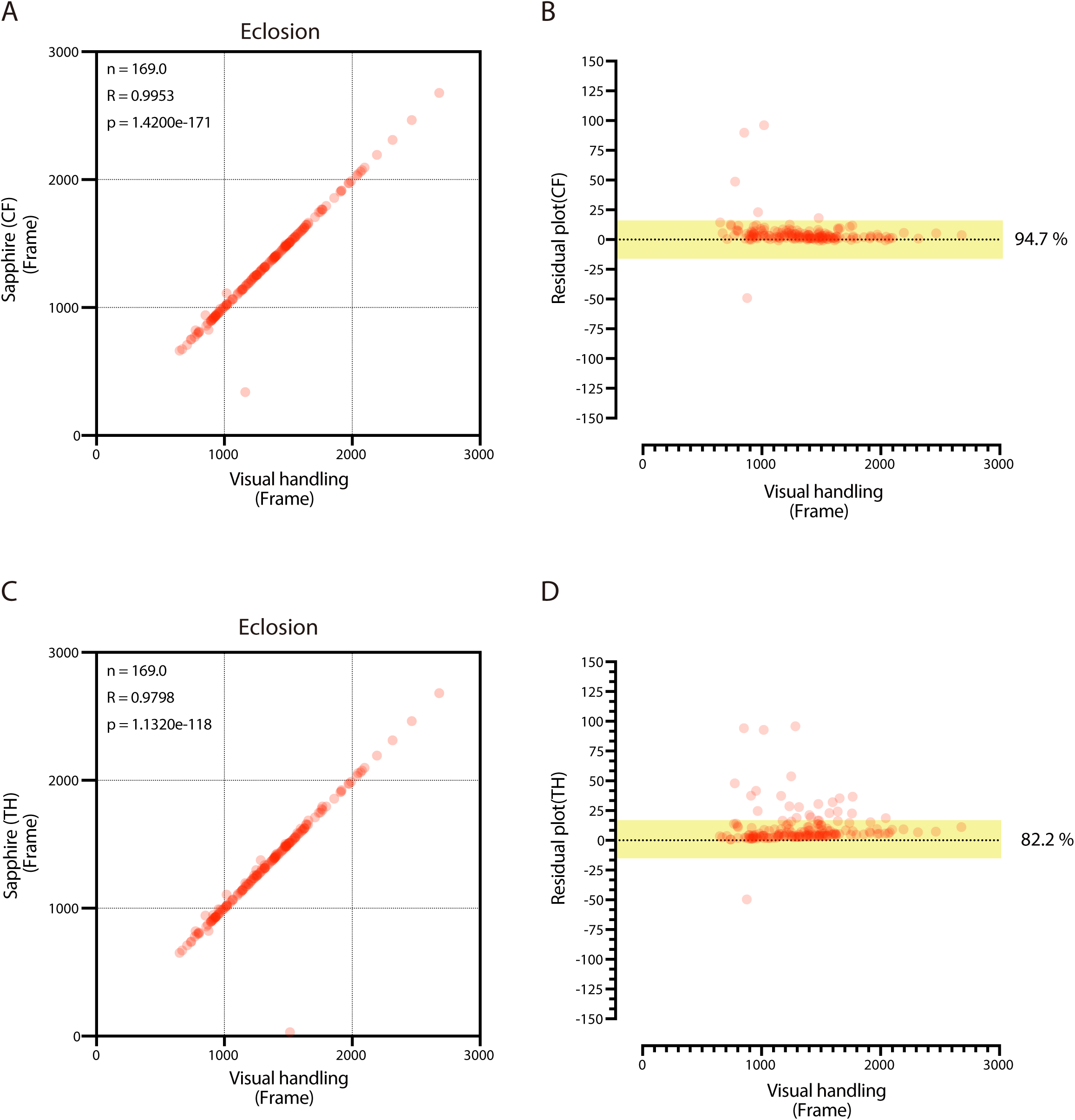
Validation of CF and TH methods. **(A-D)** Scatterplot and residual plot of eclosion timing between Sapphire (CF method: **A** and **B**; TH method: **C** and **D**) and visual handling of the same image data. Here, we transferred black pupae to new microplates once during time-lapse image acquisition.

**Figure 2-figure supplement 4.**
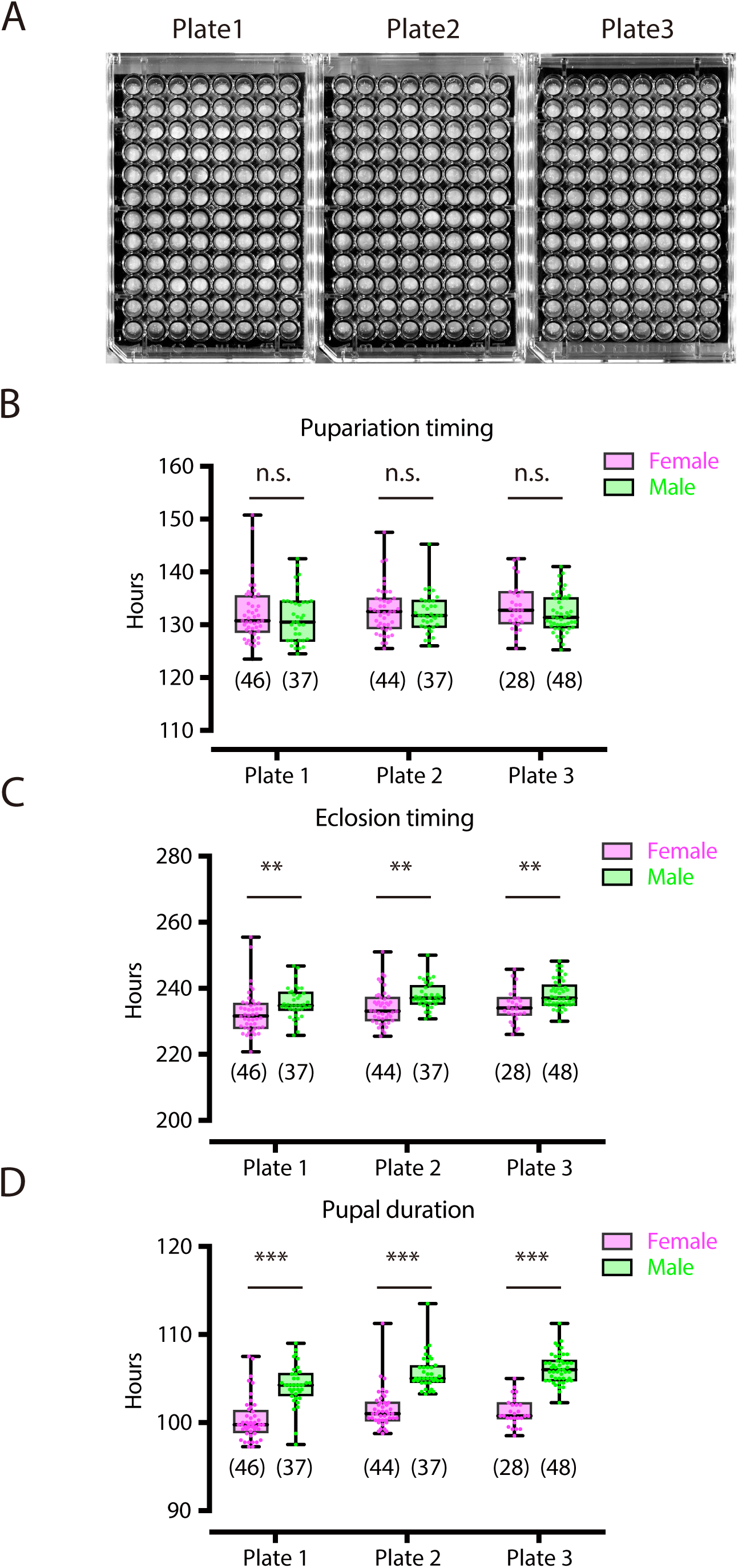
Validation of plate position effect. **(A)** Image of plates used for pupariation and eclosion time-lapse acquisition in DIAMonDS. **(B-D)** Box plots of pupariation (**B**), eclosion (**C**), and pupal duration (**D**) in male (green dots) and female (magenta dots) flies. Number of flies analyzed is indicated in parentheses on each graph. Whiskers indicate minima and maxima (\*\*\**p* < 0.001; \*\**p* < 0.01; n.s., no significant difference; Student’s unpaired *t*-test).

**Figure 3-figure supplement 1.**
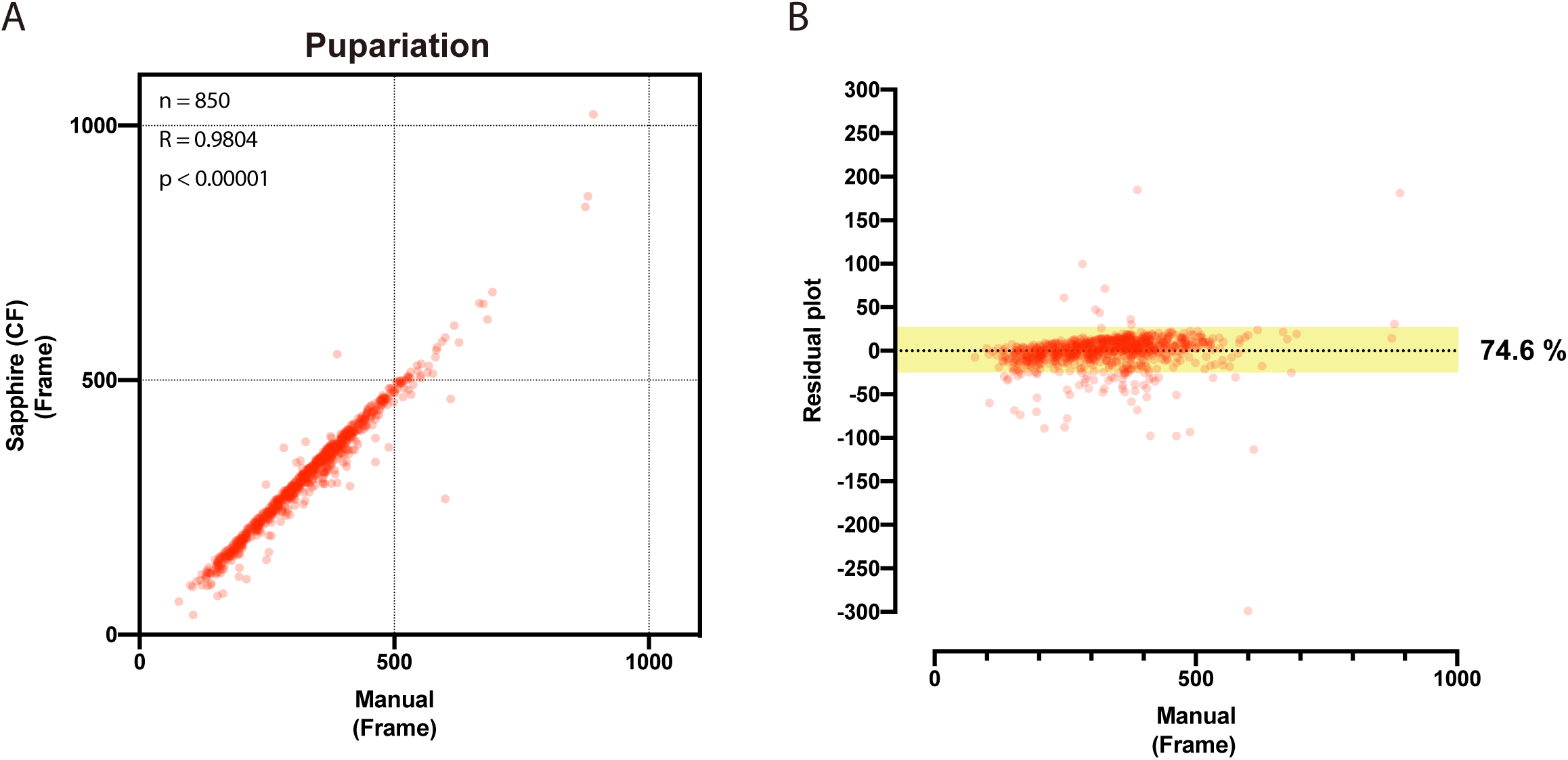
Determination of large-scale pupariation timing using DIAMonDS. **a,b** Scatterplot (**A**) and residual plot (**B**) between Sapphire (CF method) and manual analysis (n = 850).

**Figure 4-figure supplement 1.**
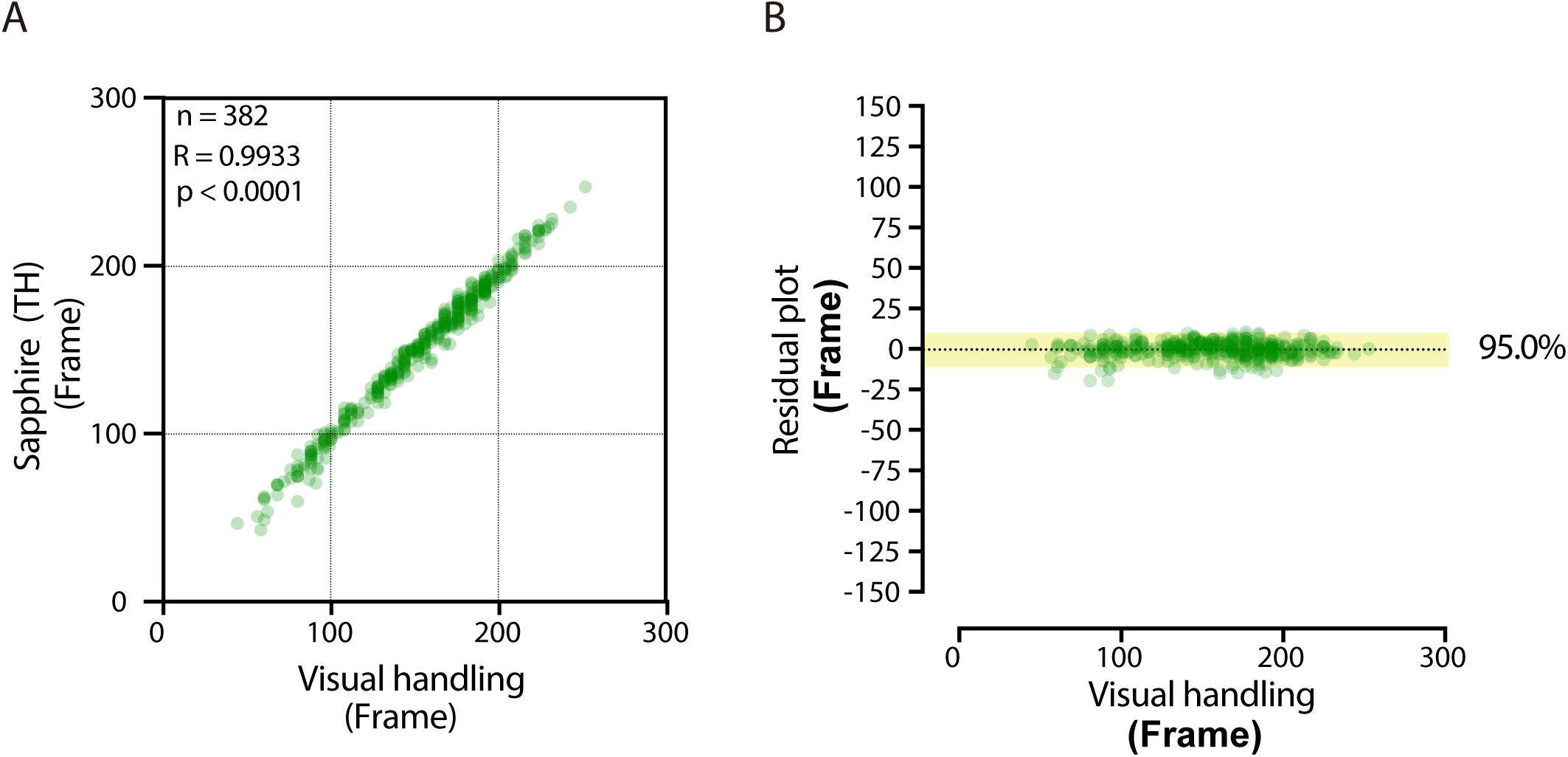
Semi-automatic TH method for Sapphire. **(A,B)** Scatterplot (**A**) and residual plot (**B**) between Sapphire (TH method) and manual analysis (visual handling) to validate accuracy (n = 382). Results corresponding to Figure. 4C,D.

**Figure 4-figure supplement 2.**
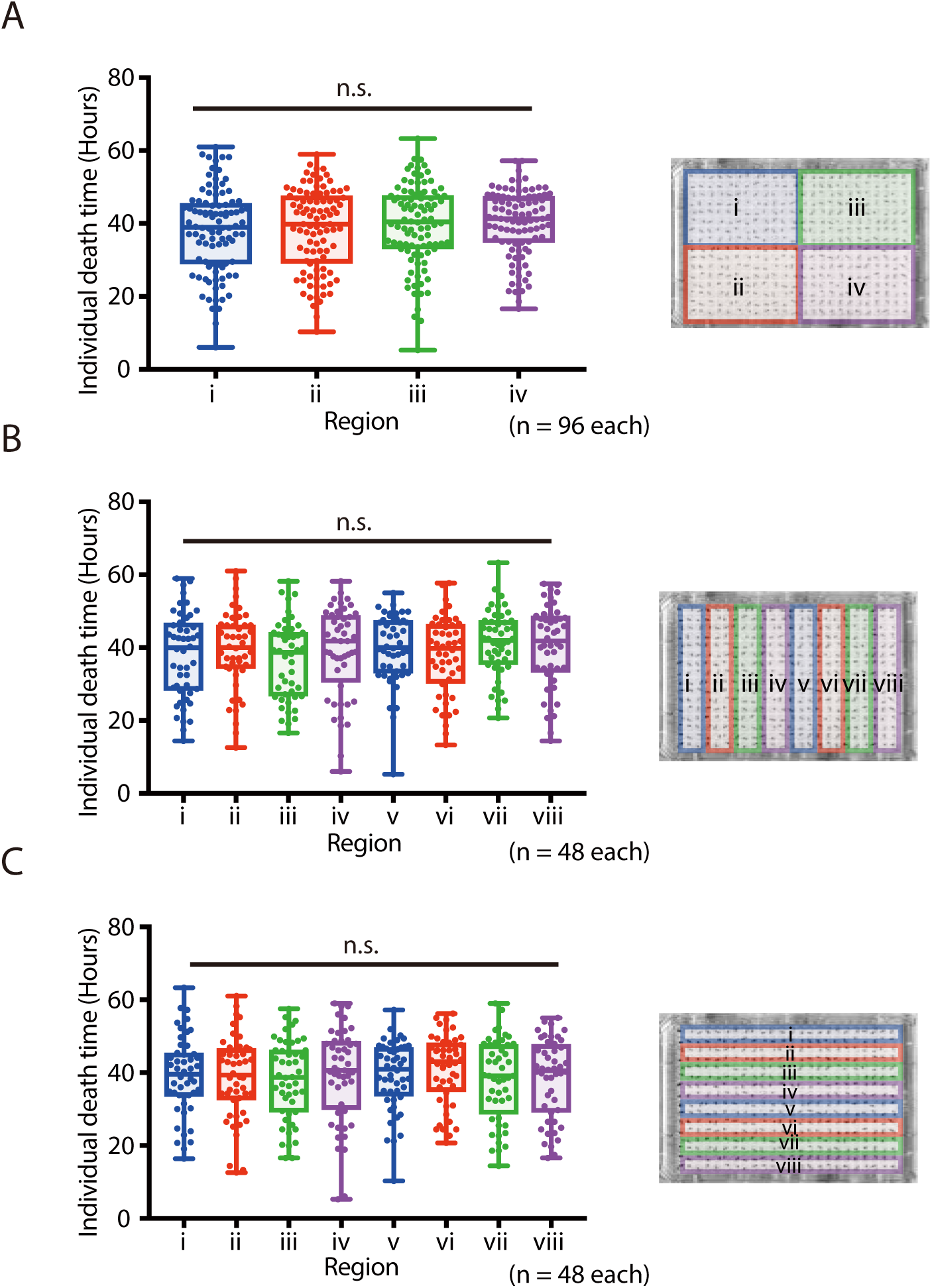
Validation of the effect of sub-area on 384-well microplate. **(A-C)** Box plots subdivided in the same dataset are shown as images on the left side of graphs. (n.s., no significant difference; Student’s unpaired *t*-test). Image of plates used to detect death timing in DIAMonDS. Number of flies analyzed indicated in parentheses on the graph.

**Figure 5-figure supplement 1.**
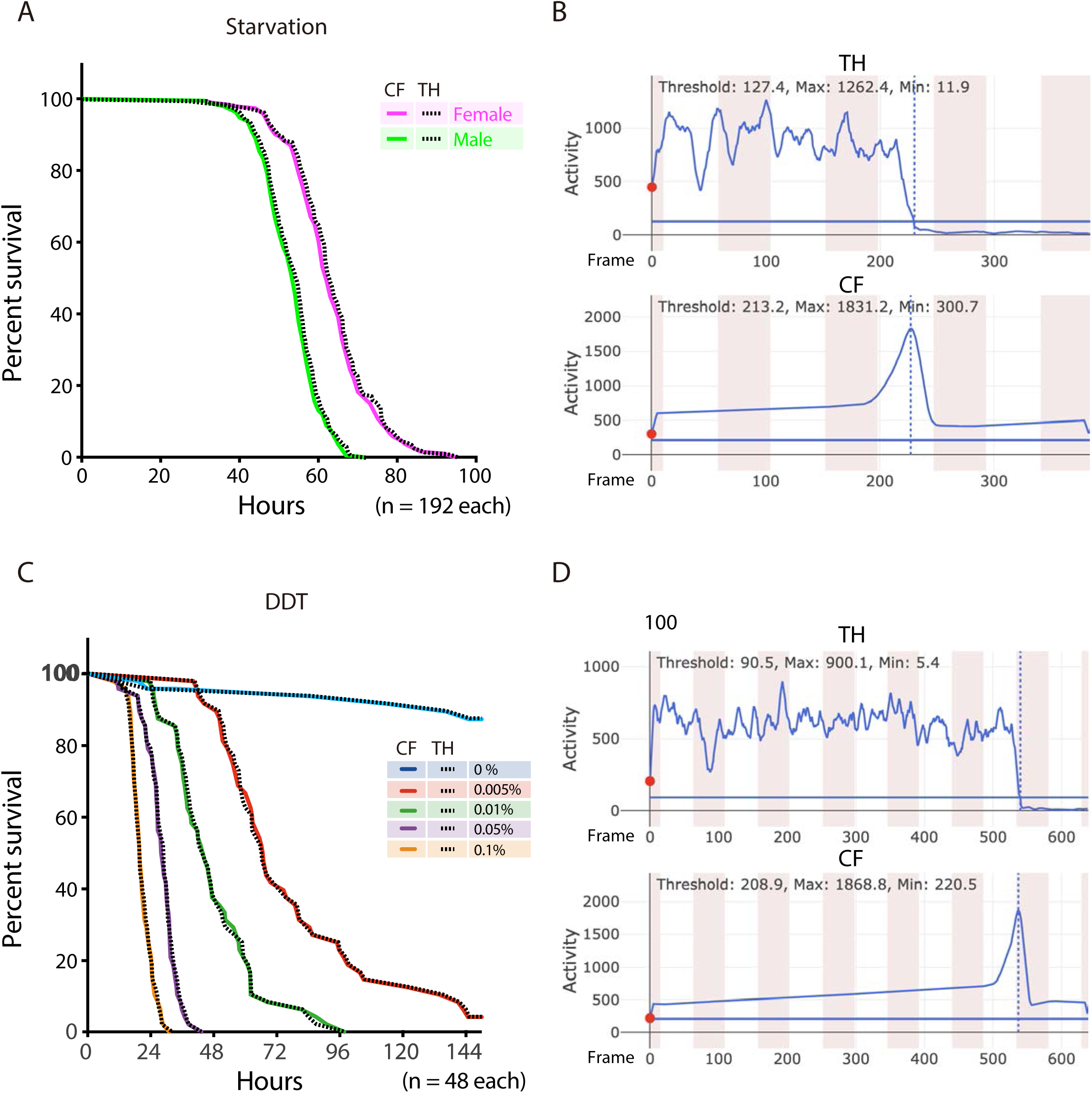
Comparison between CH and TH methods for detecting death time points in DIAMonDS. **(A,B)** Results corresponding to Figure 5A,B. **(C,D)** Results corresponding to Figure 5C,D. (**A**,**C**) Survivorship curves. Solid and dotted lines indicate results determined by CF and TH methods, respectively. (**B**,**D**) Example of analysis by Sapphire using TH and CF methods.

## Supplementary file 1 Fly Media for DIAMonDS

### 1.1. Materials and reagents

Agar powder: WAKO 010-15815 (500 g)
Agarose L (Low melting type): WAKO 317-02282 (25 g)
D(+)-Glucose: WAKO 045-31167 (10 Kg)
Sucrose: WAKO 193-00025 (500 g)
Dry yeast: Beer Yeast Korea Inc. Dry Yeast G2 (20 Kg)
Cornmeal: SUNNY MAIZE CO.,LTD. Corn grits No. 4M (25 Kg)

96-well microplate: TrueLine TR5003 96well culture plate
384-well microplate: Greiner Bio-One 781186 Clear Polystyrene 384 well Microplate
Titer stick film: WATOSON 547-KTS-HC
Fly rearing plastic vial: MKC-30 [small] (HIGH TECH LLC, Chiba, Japan)
Soft sponge plugs: COW 30t x 27⌀; (HIGH TECH LLC, Chiba, Japan)

### 1.2. Standard fly medium

-Ingredients

|  |  |
| --- | --- |
| Agar powder | 21 g |
| D(+)-Glucose | 300 g |
| Dry yeast | 150 g |
| Corn Grits | 201 g |
| Mili-Q water | 3.5 L |
| propionic acid and | 10 mL |

-Instructions

1. Measure each material and mix all.
2. stir well with heating by a table-top gas cooker until materials fully dissolved.
3. autoclave the medium.
4. Cool down the medium up to 80 °C
5. Add 10 mL propionic acid and mix well. (For fly rearing vial)
6. Dispense the medium into each vial (5 mL/vial).
7. Cool before plugging. (For preparation of DIAMonDS microplate with standard medium)
8. Dispense the medium into each well of 96-well microplate (add 170 µL for lifespan test or 100 µL for detection of pupariation and eclosion).

### 1.3. Yeast-sucrose (YS) medium for DIAMonDS drug resistance test

We have used YS medium for drug resistance test of DIAMonDS. We found that addition of 0.5% of dry yeast in 5% sucrose media showed highest survival and flies kept a high survival rate at least 10 days in the well of 384-well microplate with the food condition (Supplementary Fig. SF1.3), it seems to enough to the short-term test such as stress resistance and chemical toxicity test using 384-well microplate.

**Supplementary Fig. SF1.3.**
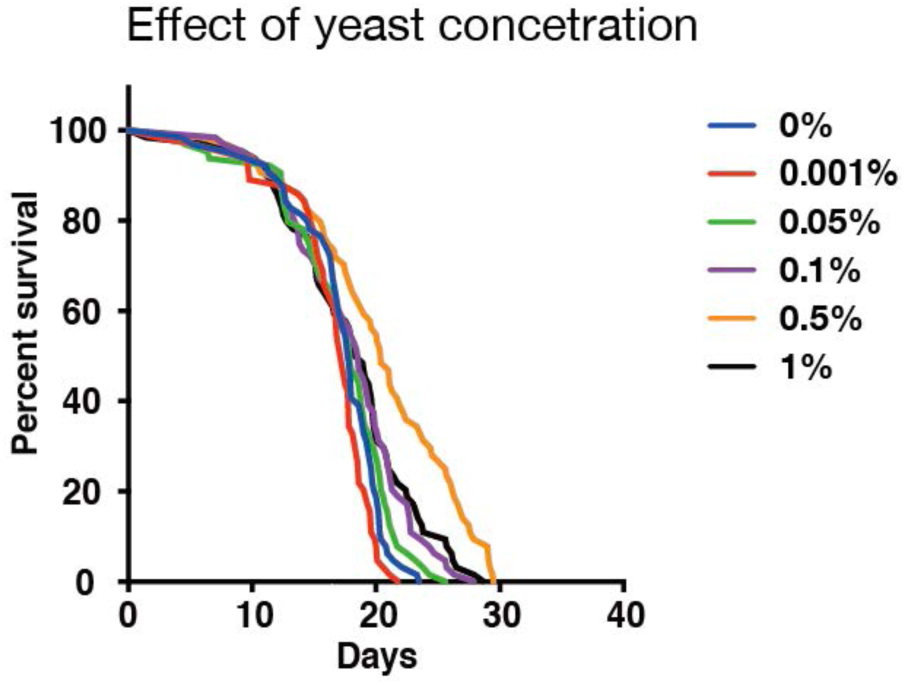
Effect of yeast concentration in fly medium.

-Ingredients

|  |  |
| --- | --- |
| Agarose L | 7.5 g |
| Sucrose | 25 g |
| Dry yeast | 2.5 g |
| Mili-Q water | (up to 500 mL) |
| propionic acid and | 1.5 mL |

-Instructions

1. Measure each material and mix all.
2. autoclave the medium.
3. Cool down and keep the medium at 50 °C by water bath.
4. Add 1.5 mL propionic acid and mix well.
5. Add appropriate volume of chemical drug and mix well.
6. Dispense the medium into each well (70 µL for 384-well and 170 µL for 96-well microplate).
7. Cool down and leave the plate until the medium surface is dry.

### 1.4. Agar plate for starvation

-Ingredients

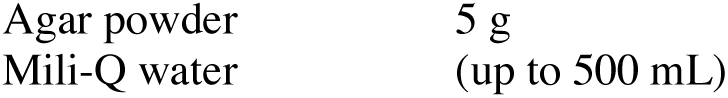

|  |  |
| --- | --- |
| Agar powder | 5 g |
| Mili-Q water | (up to 500 mL) |

-Instructions

1. Measure each material and mix all.
2. autoclave the medium.
3. Cool down the medium up to 80 °C
4. Dispense the medium into each well (70 µL for 384-well and 170 µL for 96-well microplate).
5. Cool down and leave the plate until the medium surface is dry

## Supplementary file 2 Lid for 384-well microplate

384-well microplate: Greiner Bio-One 781186 Clear Polystyrene 384 well Microplate

We have made a lid for 384-well microplate from an acrylic sheet (2 mm thick) as shown in Supplementary Fig. SF2a. Holes (diameter = 0.7 mm) were drilled in each well position with an electric drill. The four corners were fixed by screws, and all edges were sealed by masking tape (Supplementary Fig. SF2b).

**Supplementary Fig. SF2.**
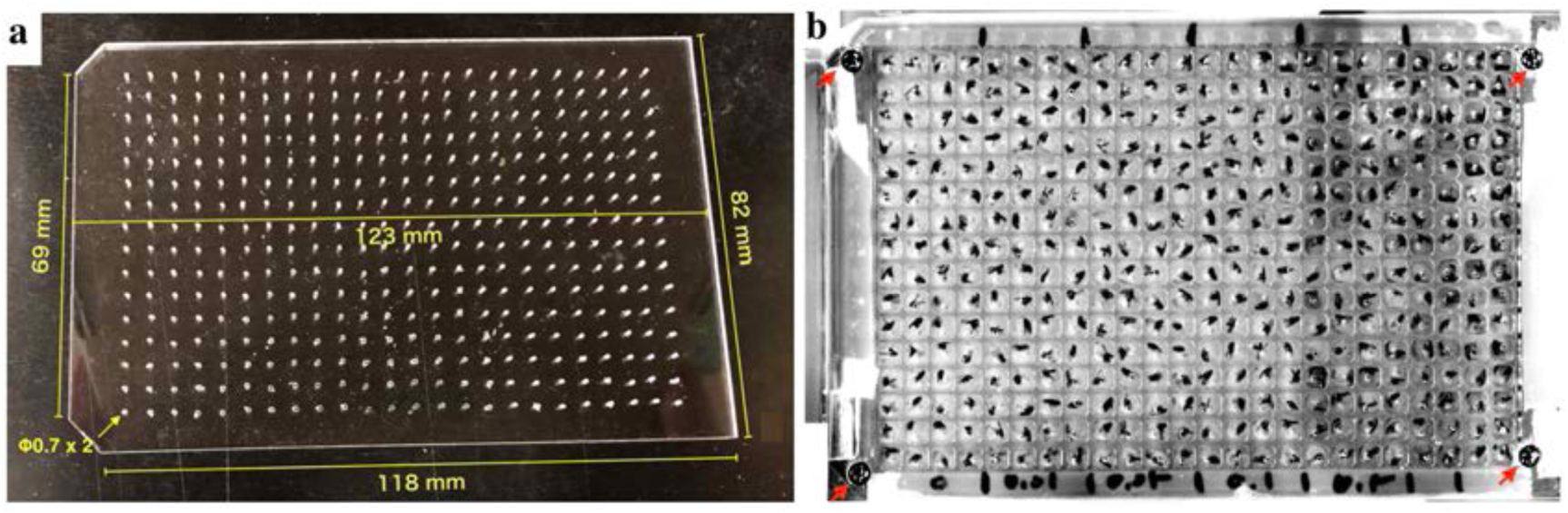
(a) A handmade acrylic lid for 384-well microplate. (b) Usage example of the lid. Red arrows indicate positions of the screw.

## Supplementary file 3 Lid for 96-well microplate

96-well microplate: TrueLine TR5003 96well culture plate
Titer stick HC film: WATOSON 547-KTS-HC

### 3-1. To measure the developmental time points of pupariation and eclosion

The titer stick film is covered on the 96-well microplate. Then, the plate is sealed on the upper surface by microplate sealing titer stick film (547-KTS-HC, WATOSON bio lab), and make eight air holes (0.35 mm) on the film surface of each well by insect pin (No. 00).

### 3-2. Stress resistance test and lifespan test

To enable transfer flies into the well in no anesthetized treatment, the cross-shaped cutting on film surfaces on each well is conducted by a sharp cutter after sealing the microplate by the film.

## Supplementary file 4 Anesthetized treatment

The use of anesthetics can reduce the labor dramatically for inserting individual flies in each well, especially in the case of a 384-well microplate. We tested two anesthetics, carbon dioxide and triethylamine (TEA, also called as FlyNap). Short exposure of CO2 (10, 30, and 60 sec) or TEA (10, 20, 30 sec) did not show a significant difference in comparison with no treated flies (Supplementary Fig. SF4a and b). The narcotism by short exposure of TEA vapor continues for 30 min^1^. Therefore, we have been preferentially using TEA for the preparation of a fly plate.

**Supplementary Fig. SF4.**
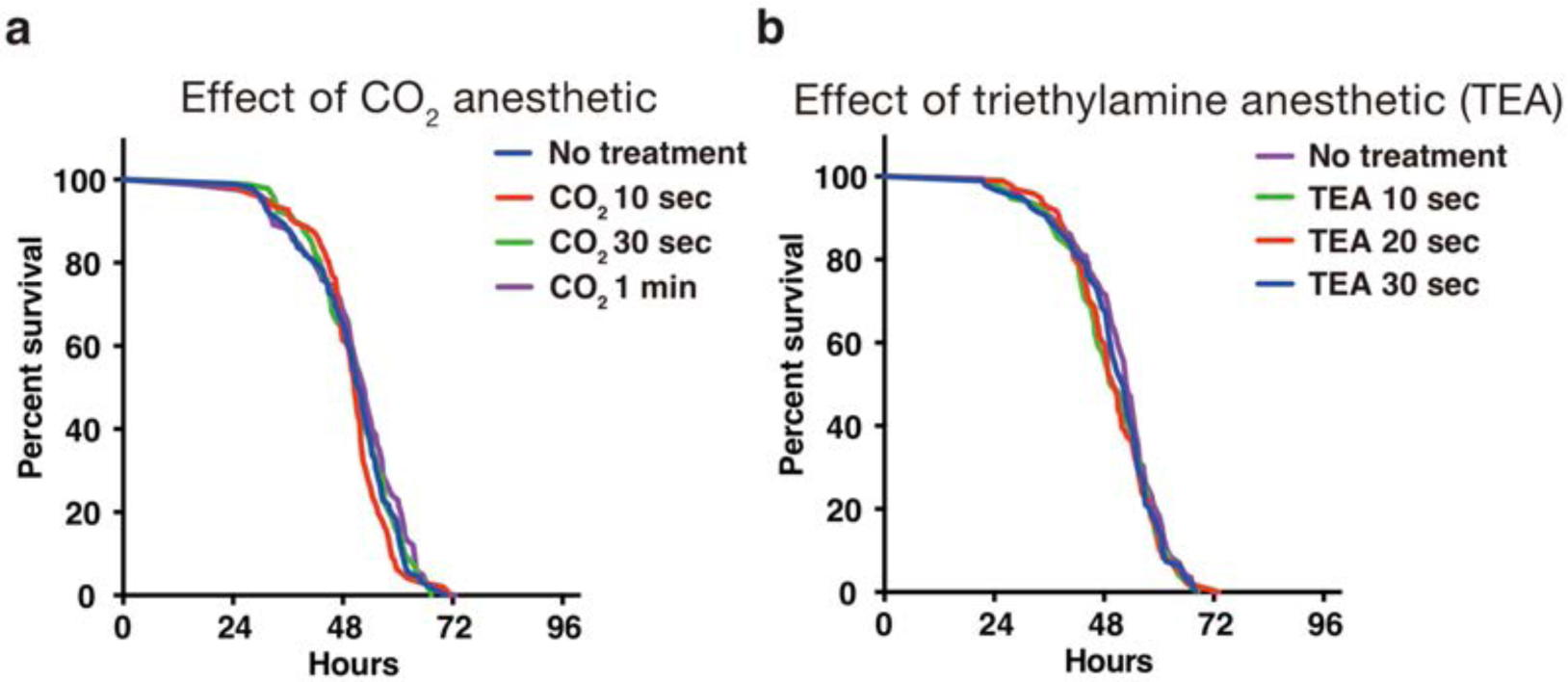
Effect of anesthetics on fly viability in starvation test of adult male flies using 384-well microplate. (a) Effect of CO2 anesthetic. (b) Effect of TEA anesthetic.

## Supplementary file 5 Fixing the microplates on the scan surface of the scanner

To put microplates on the fixed position every time, glass slides (MATSUNAMI Glass, S-2215) were arranged and glued on the scan surface as shown in Supplementary Fig. SF5a (red squares indicate positions of glass slides). The microplates were placed along the guide of slide glasses and sealed by masking tape (Supplementary Fig. SF5b and c). To reduce the reflection during scanning, black painted sheet was put on the surface of a scanner reflector (Supplementary Fig. SF5d).

**Supplementary Fig. SF5.**
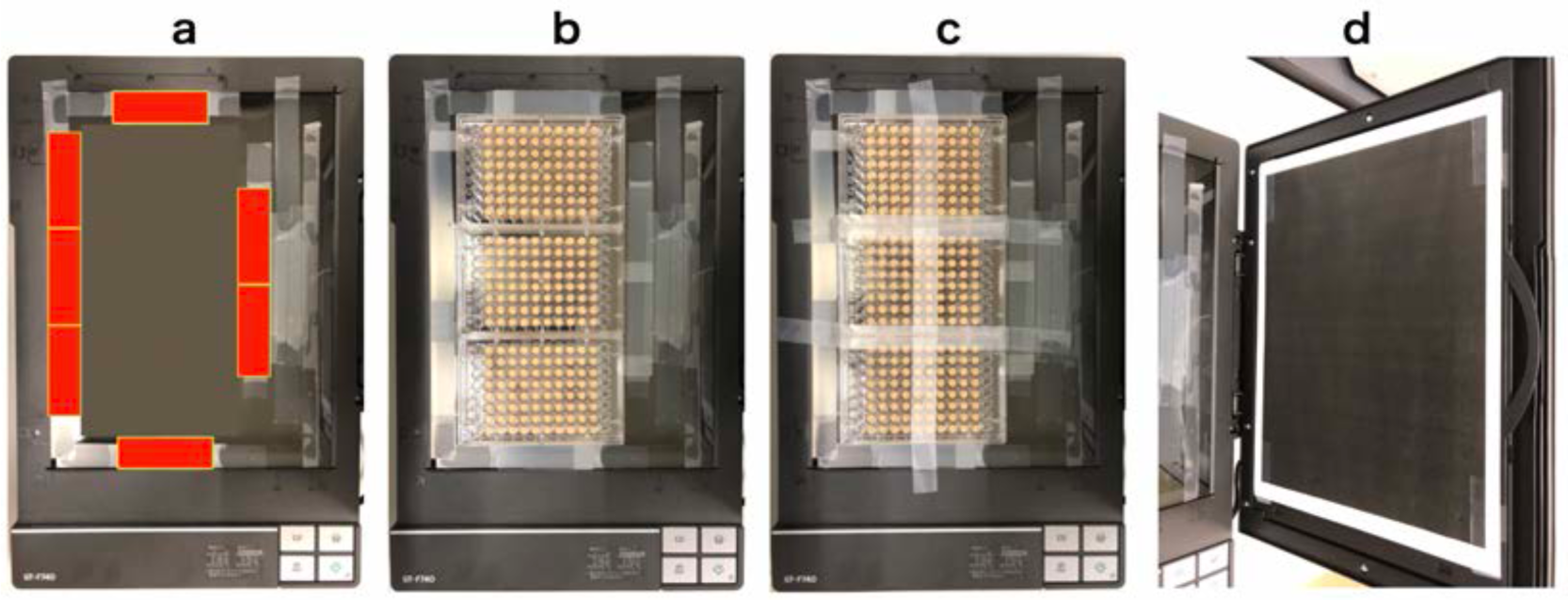
Fixing method of microplates on the scanner surface. (a) Positions of glass slides (red squares). (b) Three microplates fit perfectly in the frame of glass slides. (c) Microplates are fixed tightly on the surface by masking tape. (d) A black painted sheet on the scanner reflector.

## Supplementary file 6 Image acquisition by DIAMonDS

### 6.1. Equipments and a software

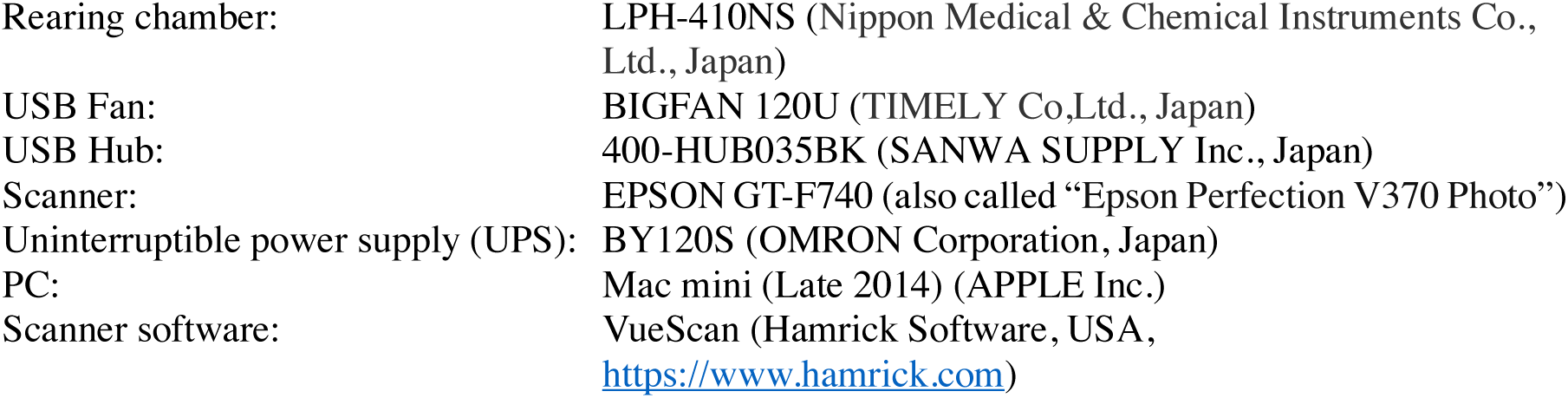

|  |  |
| --- | --- |
| Rearing chamber: | LPH-410NS (Nippon Medical & Chemical Instruments Co., Ltd., Japan) |
| USB Fan: | BIGFAN 120U (TIMELY Co.,Ltd., Japan) |
| USB Hub: | 400-HUB035BK (SANWA SUPPLY Inc., Japan) |
| Scanner: | EPSON GT-F740 (also called “Epson Perfection V370 Photo”) |
| Uninterruptible power supply (UPS): | BY120S (OMRON Corporation, Japan) |
| PC: | Mac mini (Late 2014) (APPLE Inc.) |
| Scanner software: | VueScan (Hamrick Software, USA,<br>https://www.hamrick.com) |

### 6.2. Hardware layout of the DIAMonDS

We have set up the scanner system in the rearing chamber as shown in Supplementary Fig.SF6.2a. Multiple scanners are connected to a single PC (Mac mini, late 2014, Apple Inc.). We have used an Uninterruptible power supply (UPS) (BY120S, OMRON) to reduce the risk of a blackout. To minimize temperature unevenness on the scan surface, multiple small USB-fan (d­=12 cm) are installed and operated in the chamber. DIAMonDS could use any CCD flatbed scanner. We have preferentially used EPSON GT-F740.

For the detection of individual death by DIAMonDS, the scanner is installed in the normal orientation (Supplementary Fig.SF6.2b). The medium solidified with agar gradually becomes liquified due to the feeding behavior of larvae. It is a cause of unclearness of the lid surface, and increase data noise. To reduce it, the scanner is oriented upside-down direction for measurement of developmental timings of pupariation and eclosion by DIAMonDS as shown in Supplementary Fig.SF6.2c.

**Supplementary Fig. SF6.2.**
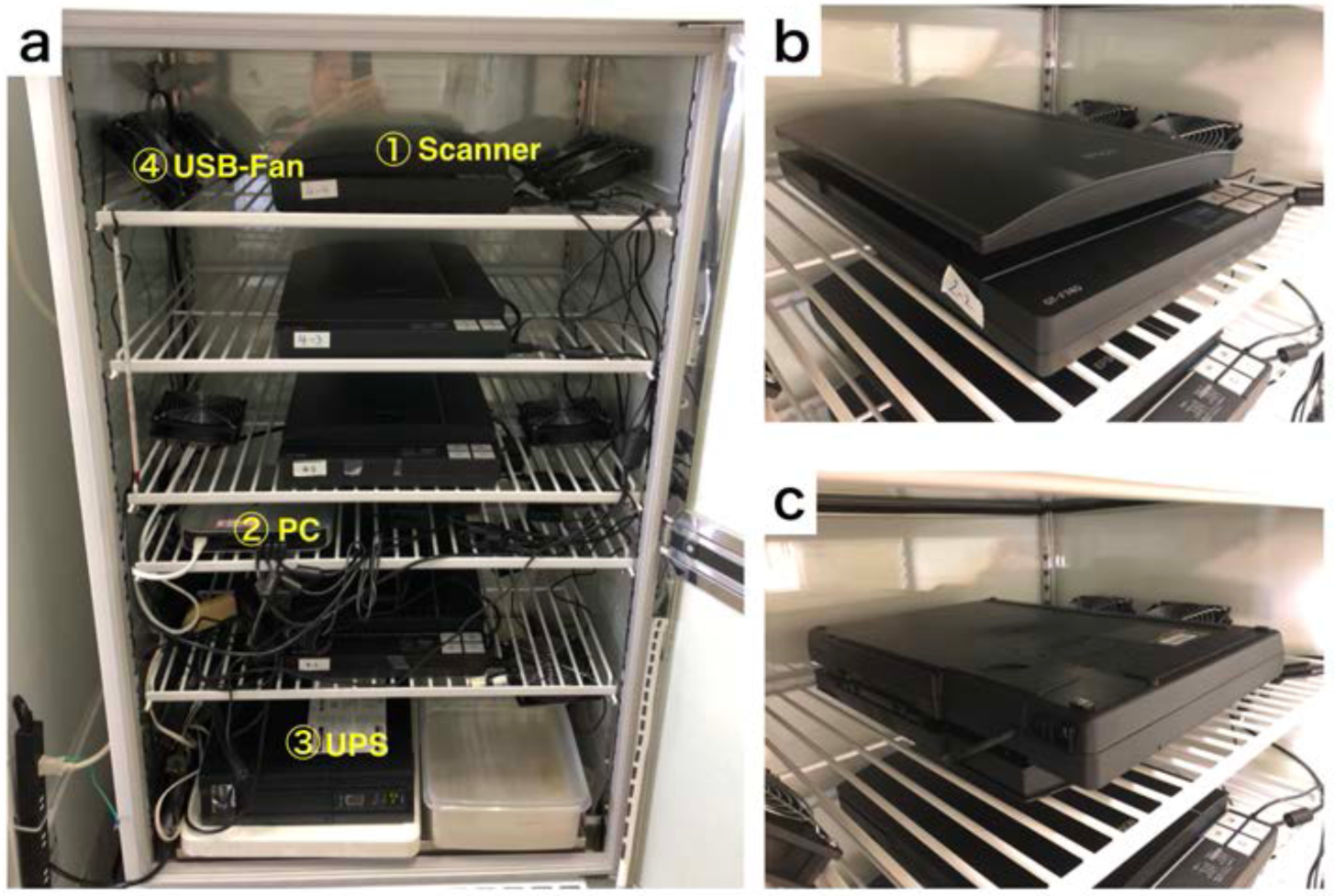
Equipment arrangement for DIAMonDs. (a) An example of the layout for DIAMonDs. (b and c) The direction of installation of the scanner for measurement of death time point. The scanner should be installed in the normal direction or upside-down direction for measurement of death (b) or developmental timings (pupariation and eclosion) (c) respectively.

### 6.3. Rearing condition

All experiments were conducted in a Plant growth chamber (LPH-410NS) which was kept at 25°C and at relative 60% humidity. Heterogeneity of temperature on the scan surface might affect data quality. To avoid the problem, several small fans (BIGFAN 120U) were installed into the chamber and the average temperature difference between several positions on scanner were achieved less than 0.5°C in dark condition (Supplementary Fig. 1).

### 6.4. Time-lapse image acquisition

To acquire the time-lapse image, we have used a scanner driver, “**VueScan**” (Hamrick Software, USA, https://www.hamrick.com). The VueScan is able to capture images continuously at several interval timing using multiple units of scanners (Smith et al. 2014). Using the VueScan, we performed time-lapse imaging as following protocol.

#### Time-lapse image acquisition protocol

1. Create new folder for storing images. An example of a folder name is “**20190707-Seong-w1118.pupariation-scanner1**” (“**date-researcher’s name-information of experiment-scanner number**”).
2. **Connect** the scanner to the PC and **turn on** the scanner.
3. Open the **VueScan**.
4. If you already saved the “**options…**”, you can load the option which recorded previously used whole parameters for acquisition of images.
5. Press the “**preview**” button, (press “**cancel**” button after completion of the preview), and set a region of interest (ROI) on the previewed image.
6. Set parameters in each tab as shown in Supplementary Fig. SF6.4.1 and SF6.4.2. In the tab “**Input**”: we use “**8 bit Gray**”, and “**150 dpi**” both as scan and preview resolutions. Press “**Auto repeat**” drop-down box and choose repeat interval as shown in Fig. SF6-4b. Select the folder to save images. Input the JPEG file name like as “**000001+.jpg**”. In the tab “**Color**”: Input the appropriate values in “**Black point (%)**”, “**white point (%)**”, “**Curve low**”, and “**Curve high**” to take a high contrast image. In the tab “**Output**”: confirm each parameter as shown in Supplementary Fig. SF6.4.2b.
7. Press **“File”/”Save options…”** and save the parameters as same folder name (e.g.: “**20190707-Seong-w1118.pupariation-scanner1**”).
8. Press “**scan**” button.
9. **(Option)** To perform multiple scanners using one PC, **repeat steps 1 to 8** for each scanner sequentially. Before the multiple scanner running, you need to replicate and assign the VueScan application for each scanner. Modify the application name of replicated each VueScan as like “VueScan-1”, “VueScan-2”… Then, open each VueScan application independently correspond to each scanner respectively.

**Supplementary Fig. SF6.4. 1.**
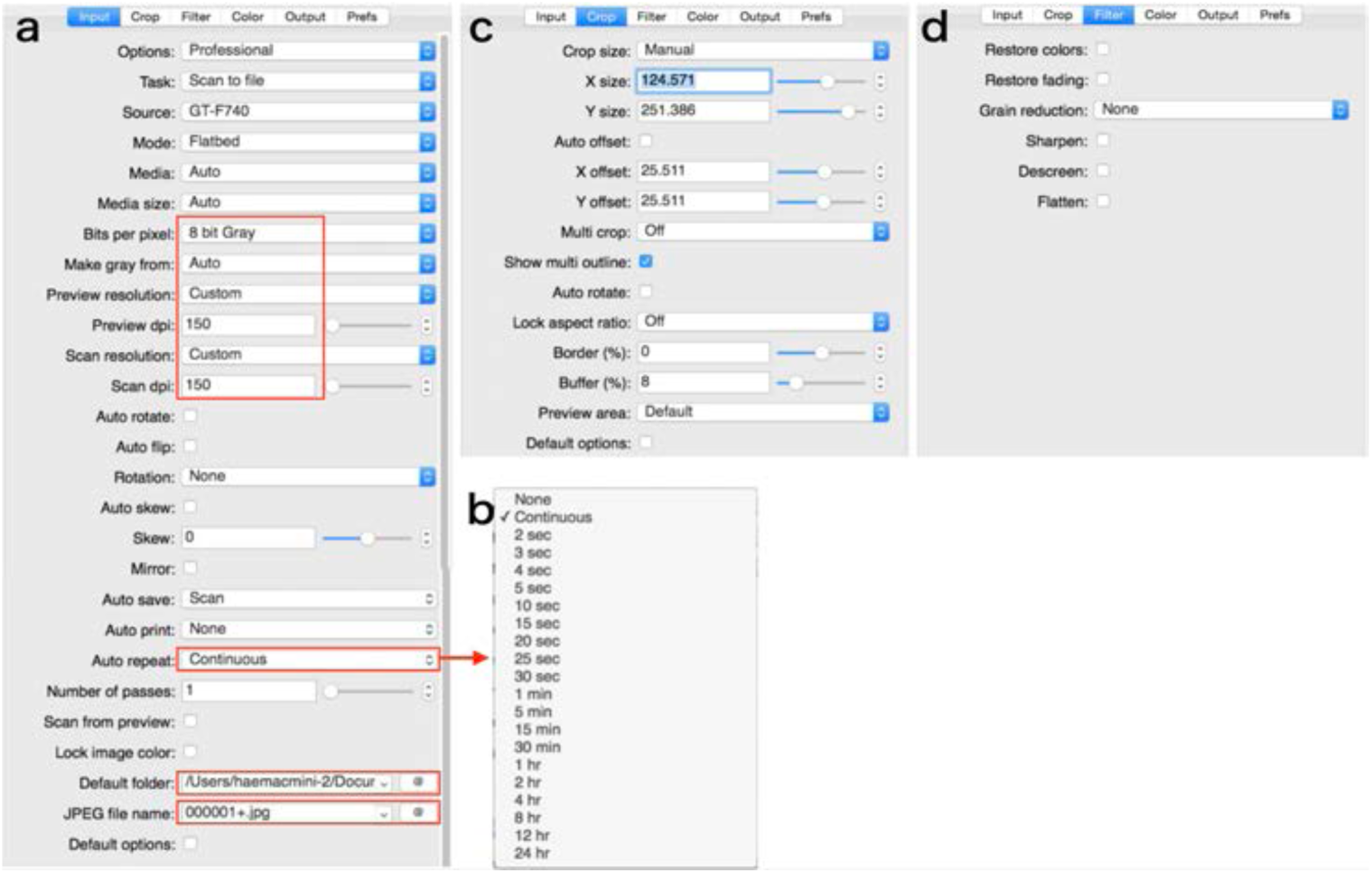
Parameters in the tabs. Areas enclosed by the square is especially important. (a) “Input” tab. (b) Drop-down box of “Auto repeat”. (c) “Crop” tab. (d) “Filter” tab.

**Supplementary Fig. SF6.4.2.**
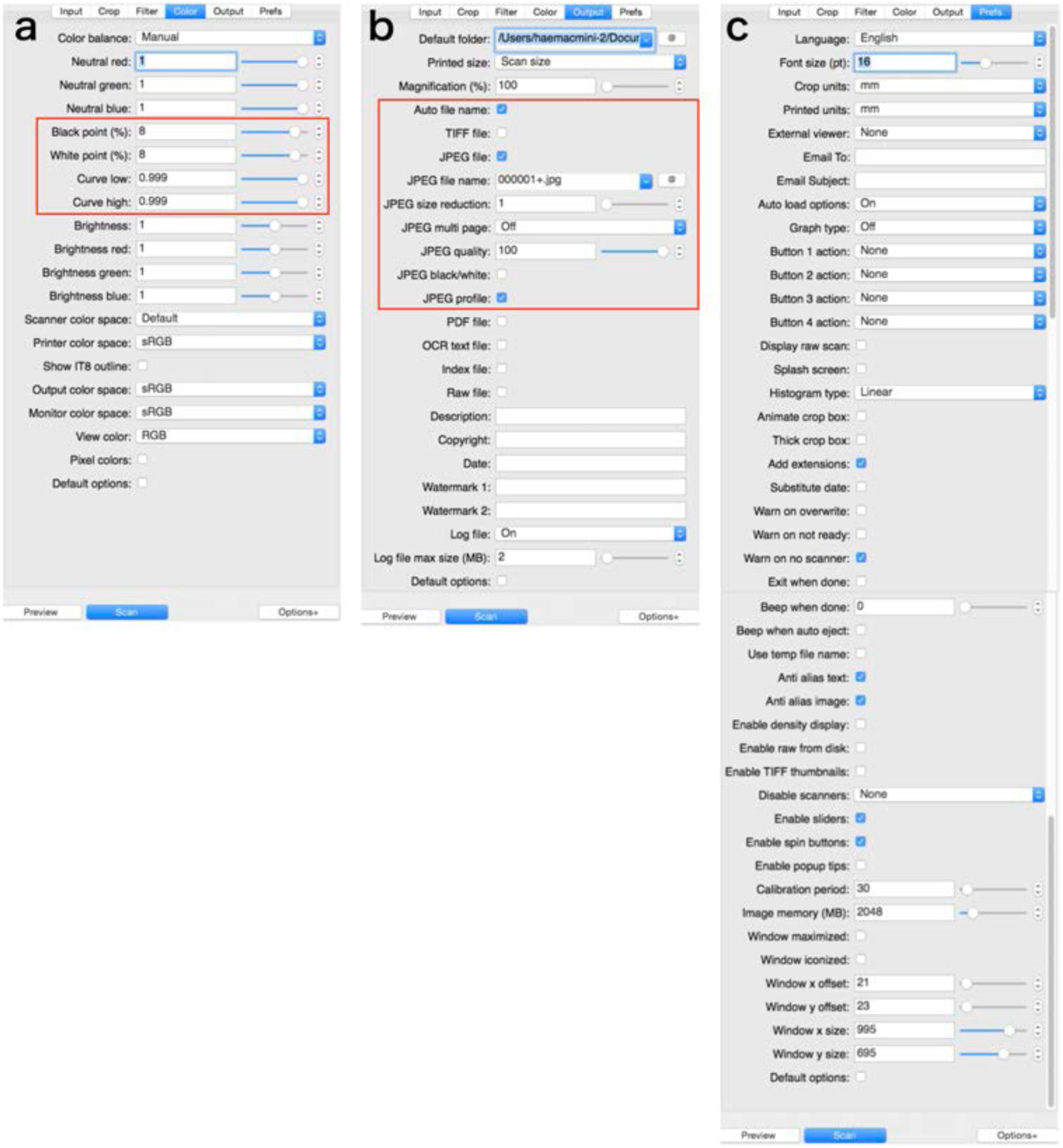
Parameters in the tabs. Areas enclosed by the square is especially important. (a) “Color” tab. (b) “Output” tab. (c) “Prefs” tab.

## Supplementary file 7 Template for analysis of DIAMonDS

We have prepared templates for data analysis of results from the Sapphire. The Sapphire analyzes the time-lapse images and, as a result, determines the event changing point as the frame number. Next, we should input each values of data (“Blacklists”, “Timestamp”, and “Event timings (auto)”) into the appropriate positions of the Excel templates. The procedure is following.

1. Download each data set (“Blacklists”, “Timestamp”, and “Event timings (auto)”) from the Sapphire viewer (Supplementary Fig. SF7.1).
  “Blacklists” contains the information of omitted wells from the analysis. We can select omitted wells in the main tab on the Sapphire viewer.
  “Timestamp” is a list of date of each image.
  “Event timings (auto)” is frame number list of each well calculated by Sapphire.
2. Copy and paste downloaded data lists into appropriate position on the template (Supplementary Fig. SF7.2 and Fig. SF7.3).
3. If necessary, input “M” or “F”. You can separate the male (“M”) and female (“F”) using the sex information of each well (Supplementary Fig. SF7.3).

**Supplementary Fig. SF7.1.**
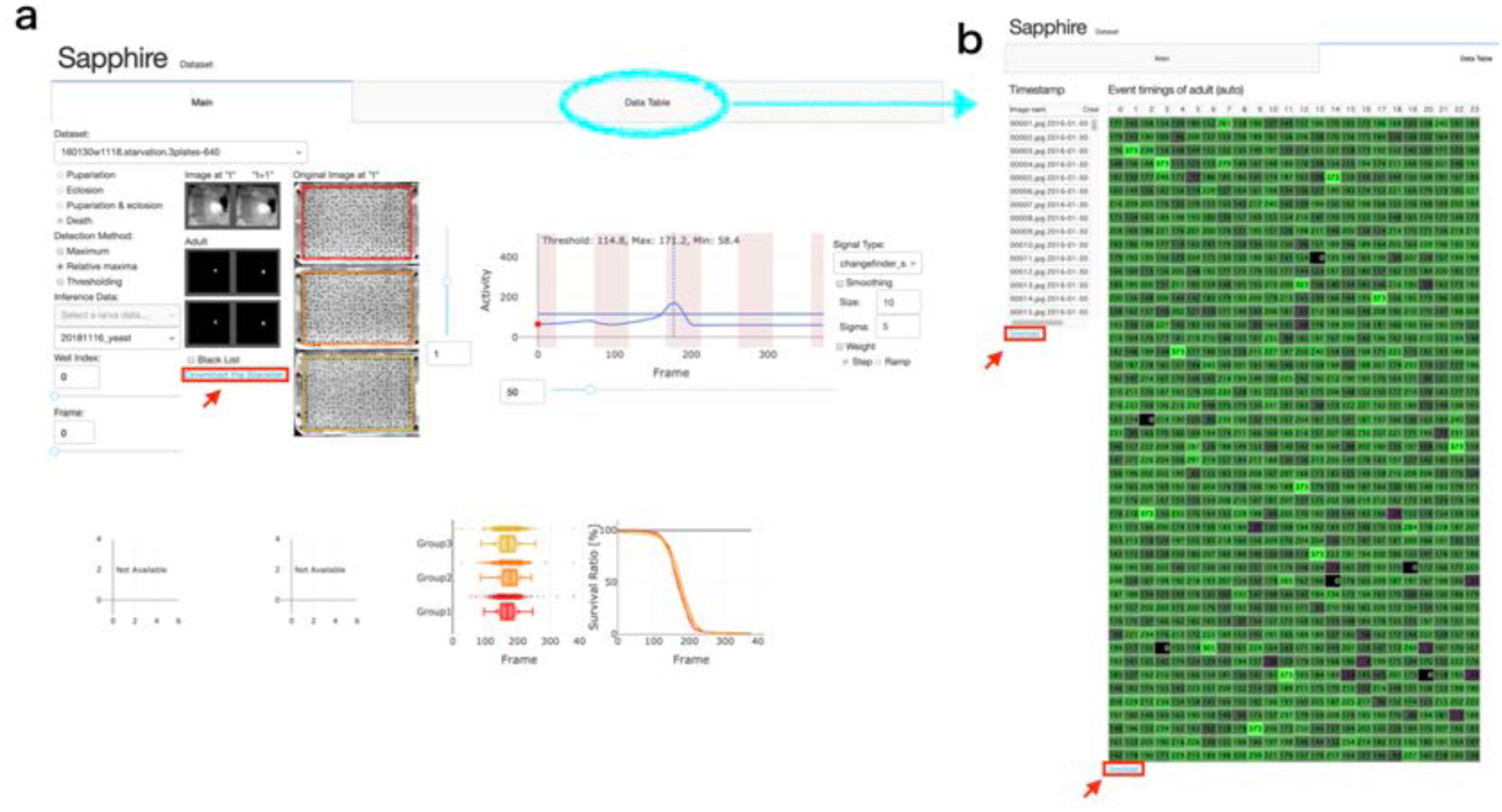
Sapphire viewer. (a) “Main” tab. (b) “Data table”. Red arrows indicated “Download” button of each data lists.

**Supplementary Fig. SF7.2.**
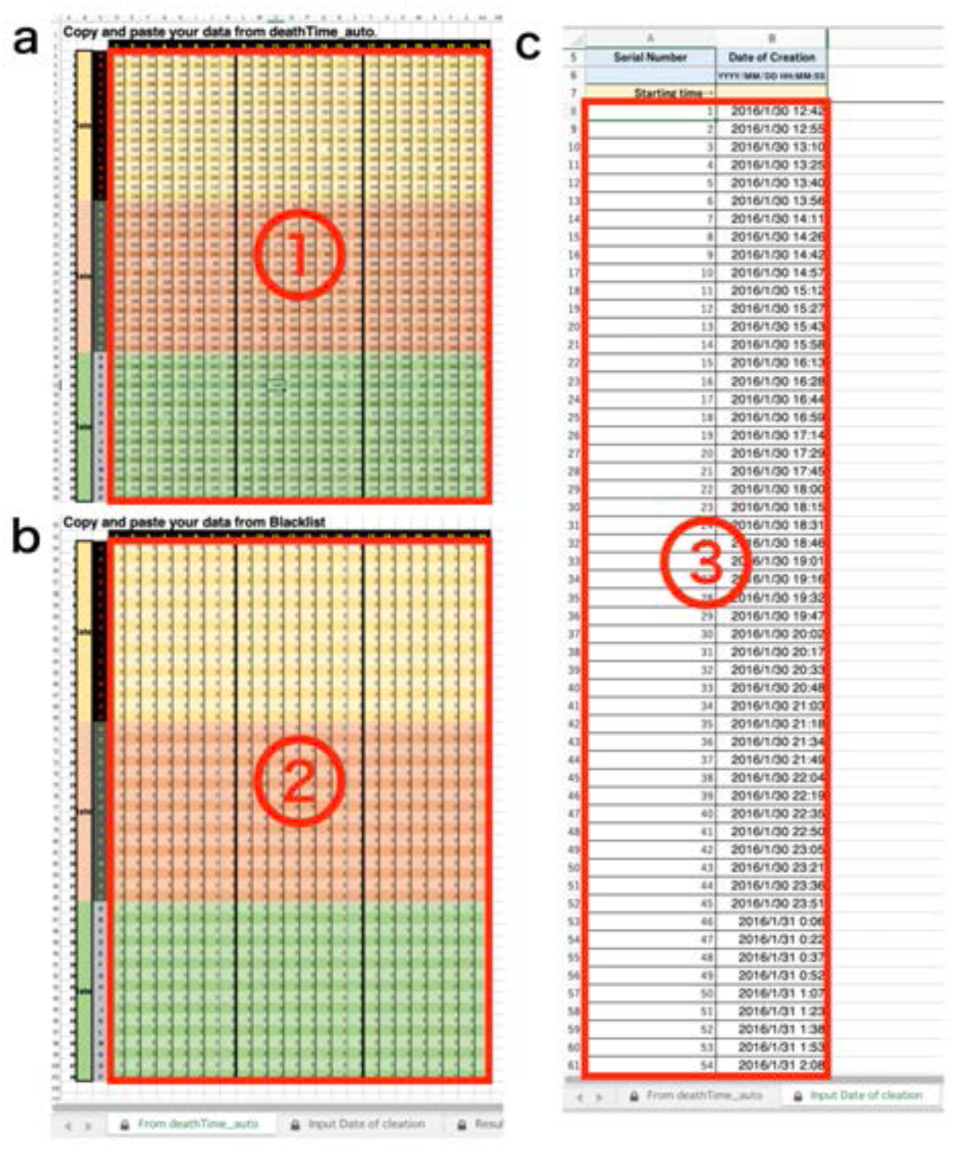
DIAMonDs analysis template: tabs for data input. (a) Input data list calculated by Sapphire into ①. (b) Input “Blacklist” downloaded from Sapphire into ②. (c) Input “Timestamp” downloaded from Sapphire into ③.

**Supplementary Fig. SF7.3.**
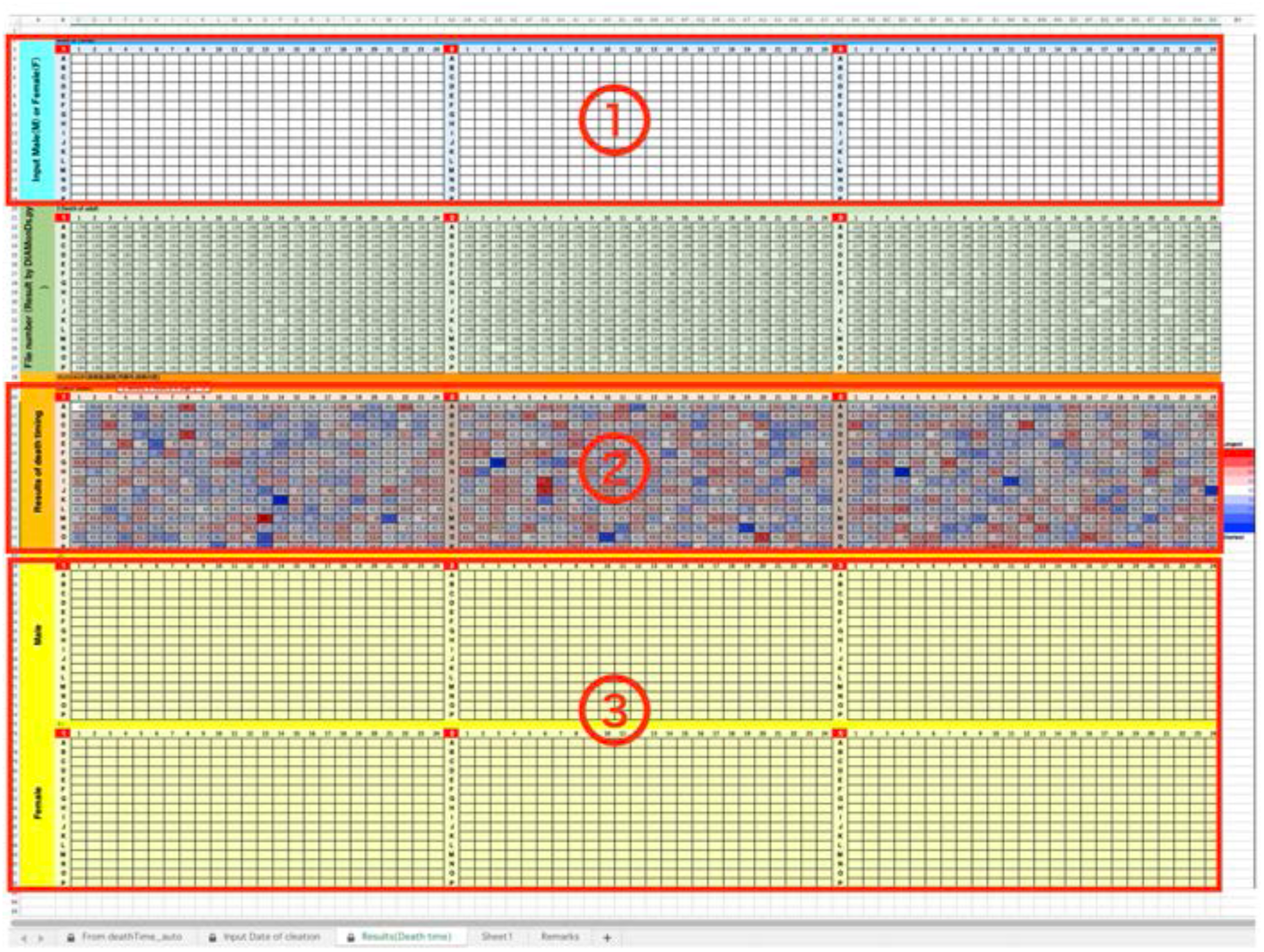
DIAMonDs analysis template: tabs for resuts. (a) Input Sex (“M” or “F”) of each well into ①. (b) The Result is indicated in ②. (c) In ③, Results are separated two groups, male and female.

## Supplementary file 8 Life-event detection algorithm and software: Sapphire

Present system provides automatic high-accurate life-event detection algorithm including the image processing and signal processing shown in Figure 1c. Following text describes quantitative summary and robustness of the algorithm in various parameter spaces.

### 8.1. Data augmentation for network training

**Supplementary Table SF8.1.**
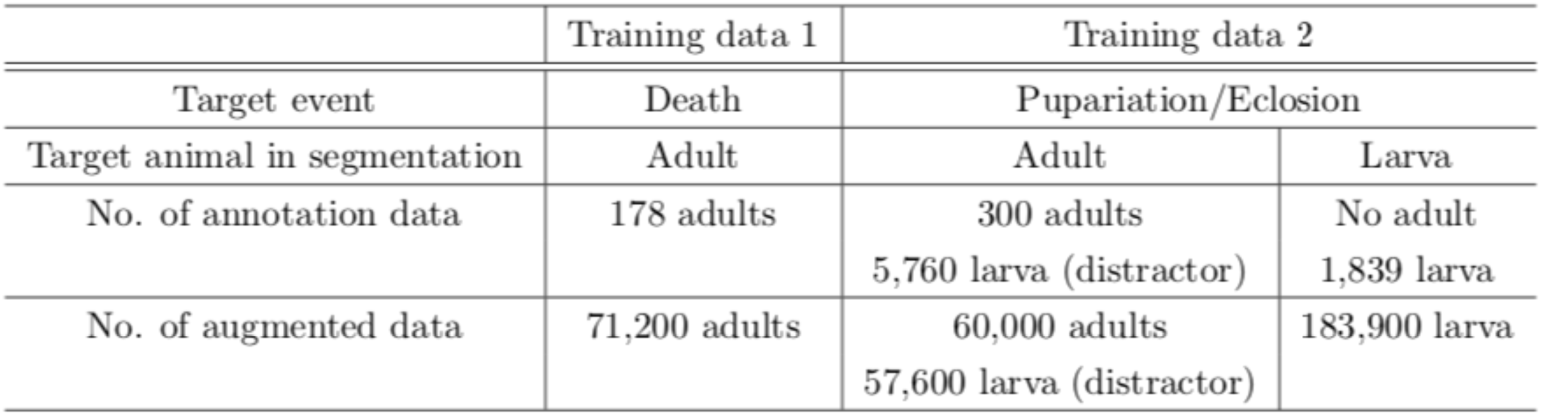
Data augmentation for training of FCN designed for animal body segmentation. Note that the system was trained by only two datasets.

### 8.2. Animal body segmentation

Present system could detect rough outline of animal body (Supplementary file 8A).

In some cases, segmentation errors still appeared. The errors in segmentation could be improved by the increase of annotation data for training of network. In other word, present system showed high performance even though datasets shown in the manuscript are obtained through several imaging condition and different laboratories as well as the number of annotation data was small. In consistent imaging condition (same shape and size of well, plate number, and so on), the system will robustly detect entire life-event. Additional annotation data for every experiment might be also effective in order to promote network training. In other word, present algorithm could reach sufficient accuracy if an appropriate signal processing was applied on the image data even with rough segmentation.

### 8.3. Robustness of detection for CF parameters

In the present system, subtraction between consecutive segmentation images were converted to ChangeFinder (CF) signal which is one of change point detection algorithms^1^ (see *online method*).

For the death detection, the system calculates the CF signal from adult body segmentation (Supplementary file 8Ba). For the pupariation and eclosion detections, two CF signals were calculated from adult and larva segmentations, respectively (Supplementary file 8Bb).

In the present system, the timing of life-event transitions is determined as the maximum point of corresponding CF scores (‘Max-CF’).

To evaluate robustness of detection, we compared several detection methods. Most simple way is just thresholding of the activity signal without change point detection such as CF. We calculated two types of signal from difference between consecutive images. One is the luminance difference (‘Luminance diff.’) obtained by subtraction between raw images, and another one is the subtraction between segmentation images before the CF (‘Seg. diff.’). Threshold for event timing were calculated as total average of the signals (‘auto’) and manual tuning (‘manual’). In comparison of these event detection methods, max-CF exhibited high accuracy in almost all datasets (Supplementary Table SF8.2).

**Supplementary Table SF8.2.**
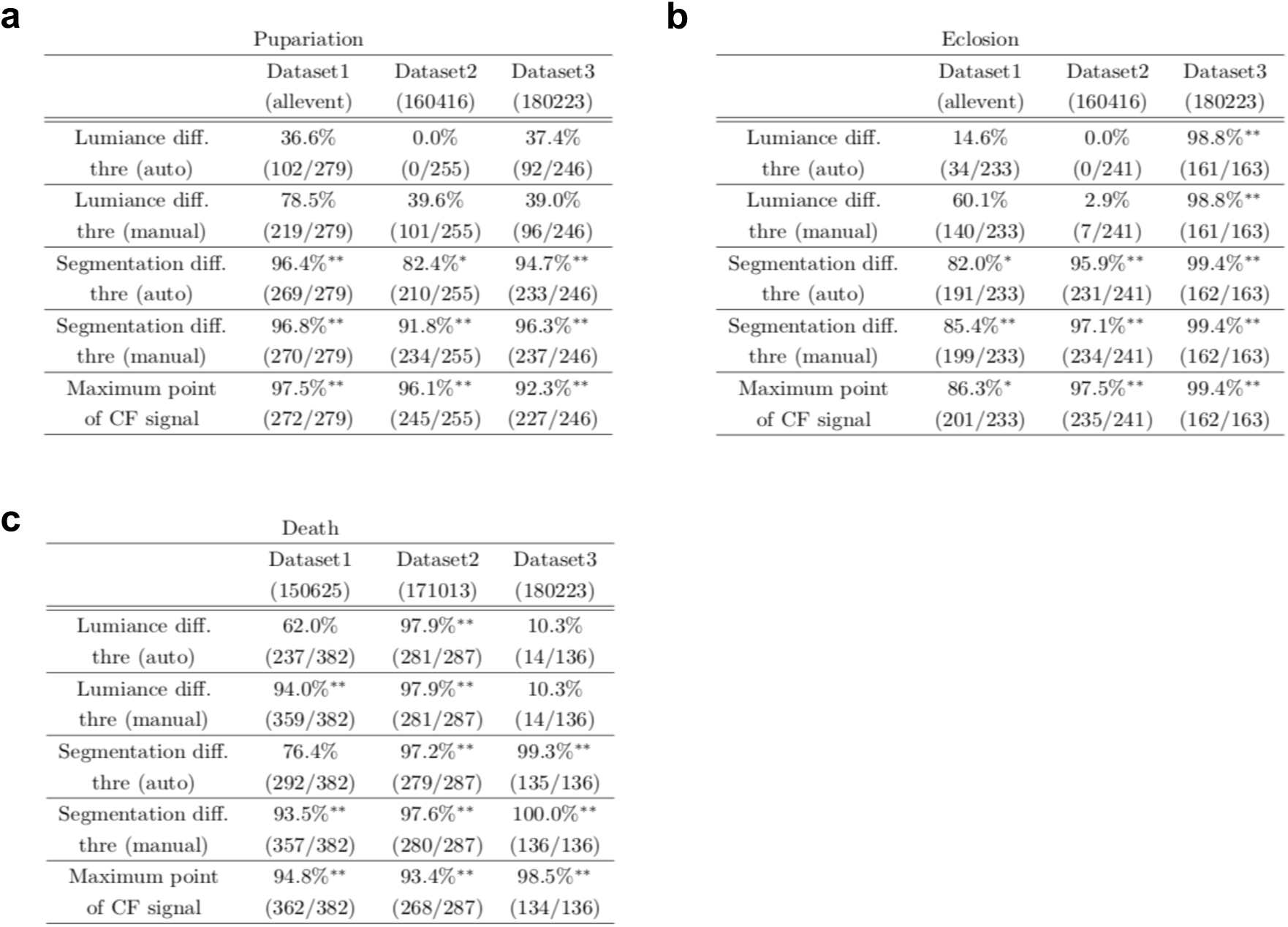
Summary of consistency between automatic and manual detections of pupariation (a), eclosion (b), and death (c). The consistency was evaluated as the ratio of individuals whose detected frame differences between the algorithm and human expert were less than 5% for entire frame. Columns indicates different dataset and rows corresponds different detection method for comparison. Consistencies were described as the ratio and the number. The cells with >=90% consistency were labeled by two asterisks, and one asterisk corresponds the cells equal or greater than 80%, and less than 90%. Note that the total numbers of animal were different with respect to dataset because of the elimination of some individuals that could not had the events.

Supplementary Table SF8.2 Summary of consistency between automatic and manual detections of Pupariation (a), Eclosion (b), and Death (c). The consistency was evaluated as the ratio of individuals whose detected frame differences between the algorithm and human expert were less than 5% for entire

CF has two parameters, *r* and T. The *r* is a discounting rate and *T* is the length of a time window for averaging. We examined in robustness of detection accuracy for these two parameters in max-CF. For all event target and all datasets, max-CF demonstrated high-accuracy in wide range of the parameter, even the accuracy is relatively sensitive for discounting *r* than smoothing *T* (Supplementary Fig. SF8.3.1-SF8.3.3).

**Supplementary Fig. SF8.3.1.**
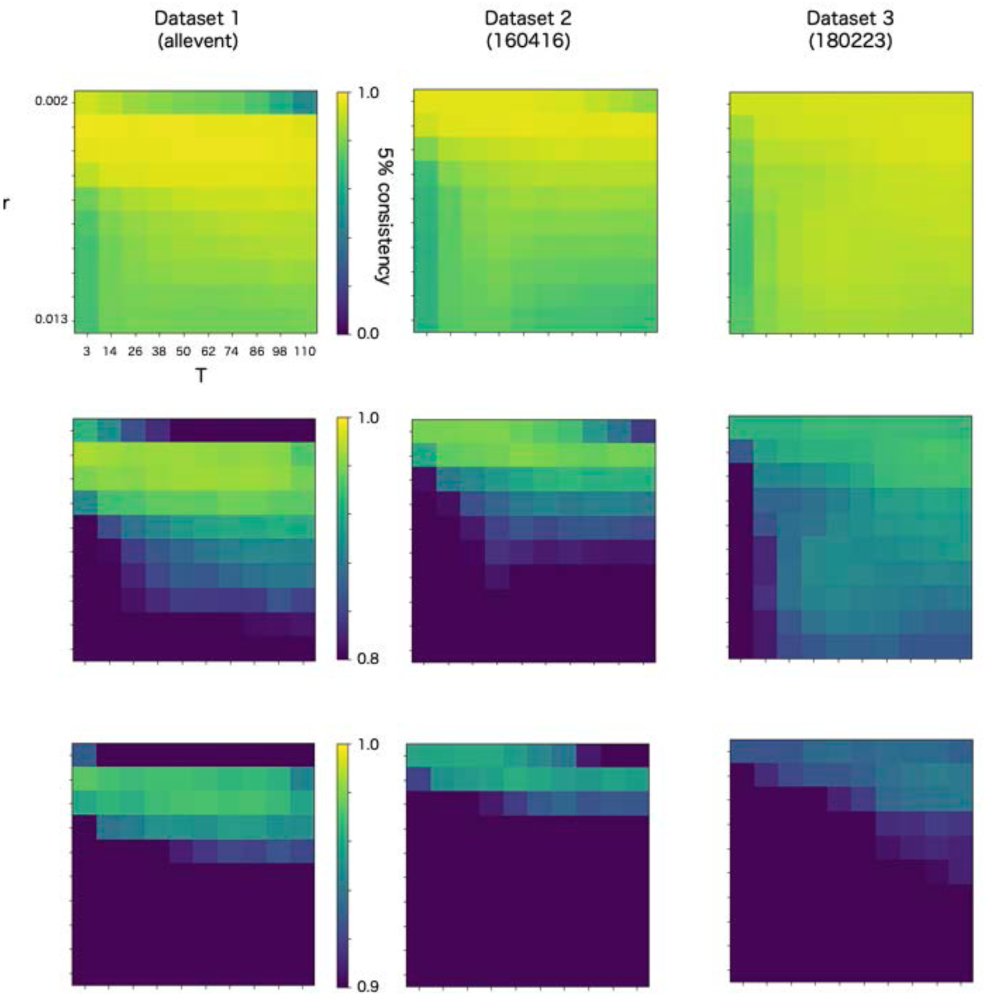
Robustness of automatic detection system for ChangeFinder parameters at pupariation detection. Each panel shows 5% consistency as heatmap for *T* (x-axis) and *r* (y-axis) of ChangeFinder’s parameter. Panels are placed for different datasets (column) and color-scales (row).

**Supplementary Fig. SF8.3.2.**
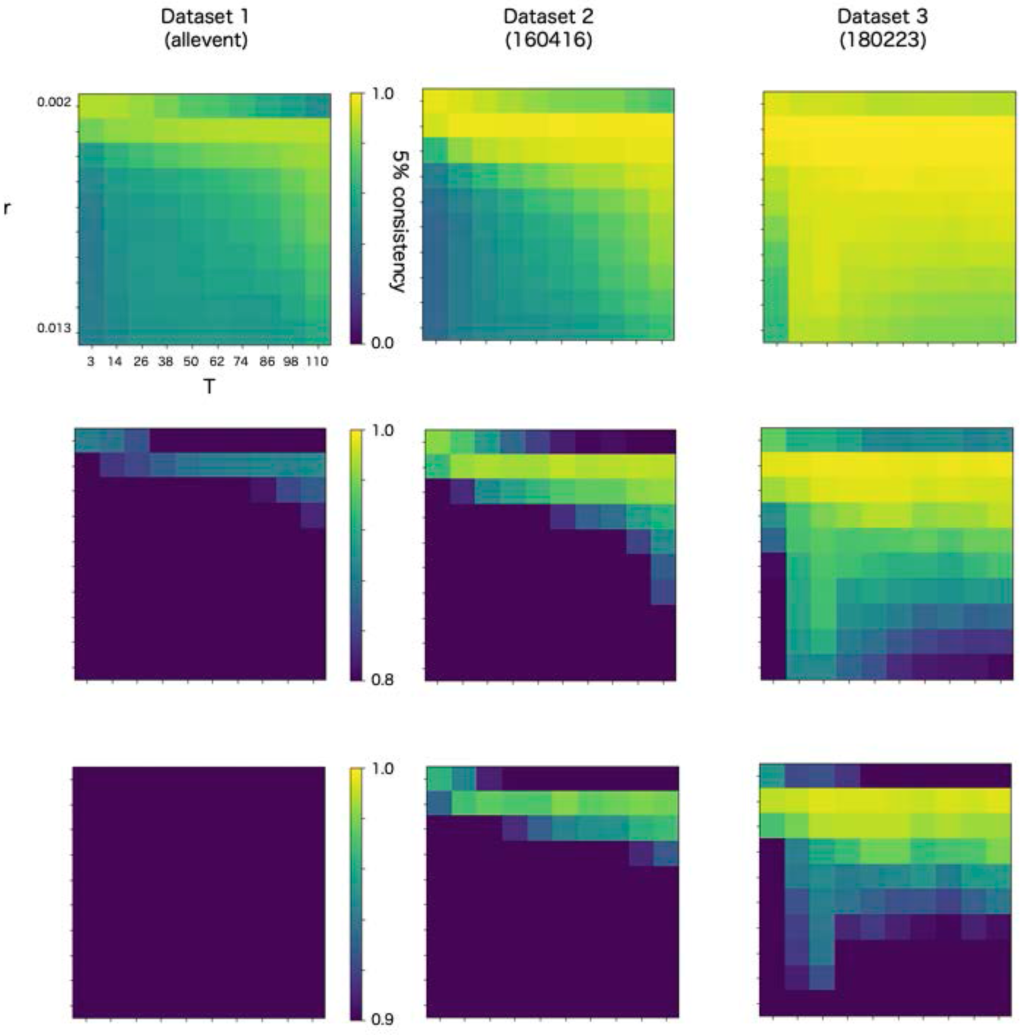
Robustness of automatic detection system for ChangeFinder parameters at eclosion detection. Each panel shows 5% consistency as heatmap for *T* (x-axis) and *r* (y-axis) of ChangeFinder’s parameter. Panels are placed for different datasets (column) and color-scales (row).

**Supplementary Fig. SF8.3.3.**
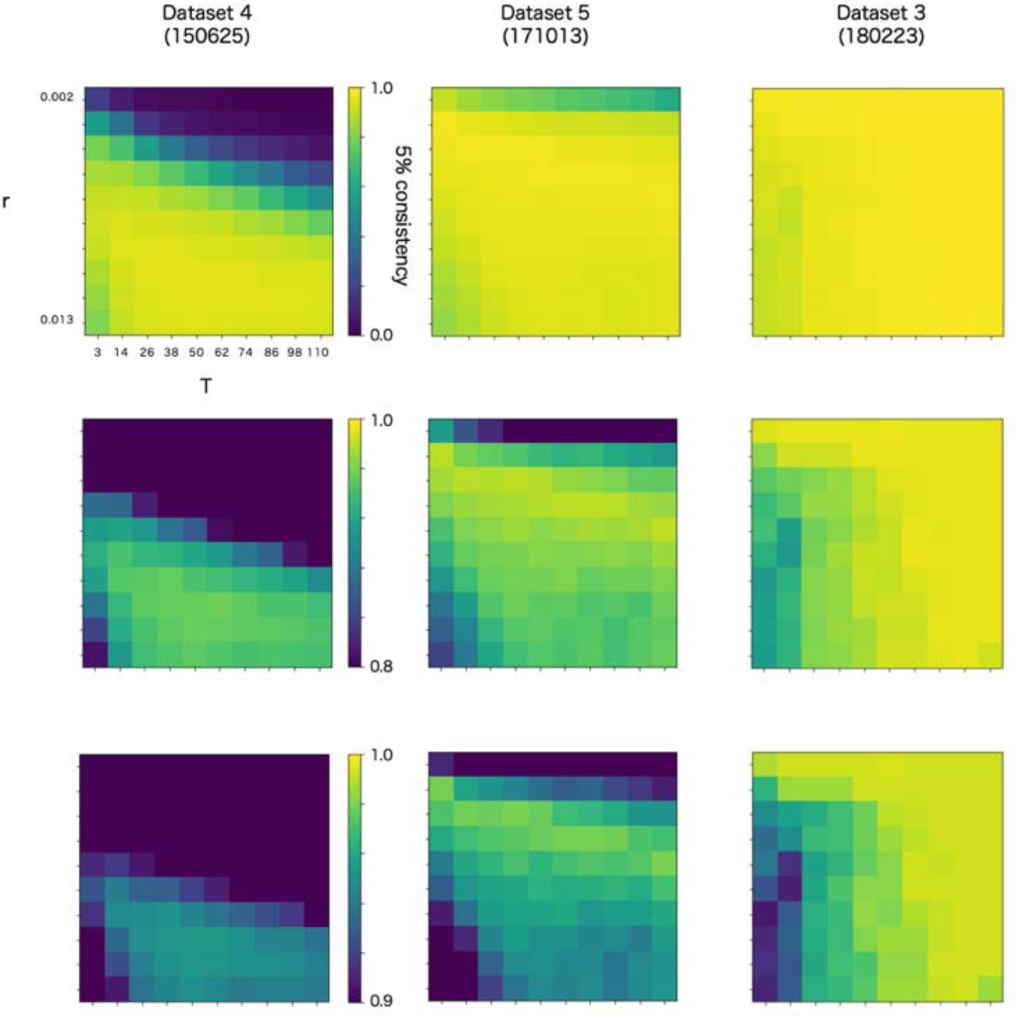
Robustness of automatic detection system for ChangeFinder parameters at death detection. Each panel shows 5% consistency as heatmap for *T* (x-axis) and *r* (y-axis) of ChangeFinder’s parameter. Panels are placed for different datasets (column) and color-scales (row). Most right column (dataset 3, 180223) is same dataset in the Supplementary Fig. SF8.3.1 and SF8.3.2.

Present system used *r*=0.003 except death detection in the dataset 1 (150626). In the dataset, *r*=0.009 because the dataset has significantly few images.

### 8.4. Application of the algorithm

The system captures the animal body and its trajectory. Therefore, one can estimate individual activity for entire recording as the summation of the trajectory. (Supplementary Video 1). In addition, the system can demonstrate dynamic visualization of life-event transitions of drosophila population (Supplementary Video 2).

---

**Supplementary Video 1.**
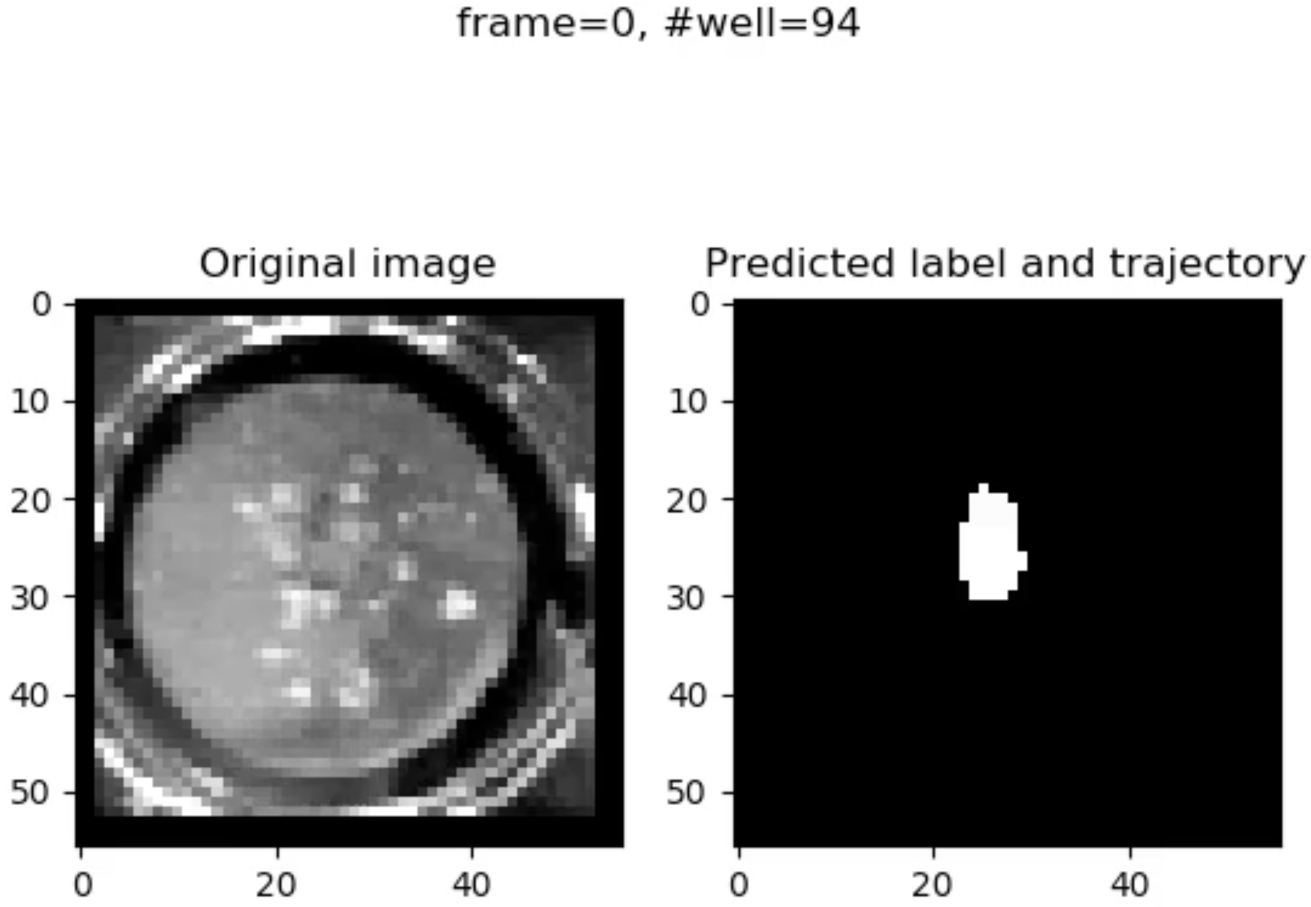
Trajectory of individual animal movement.

**Supplementary Video 2.**
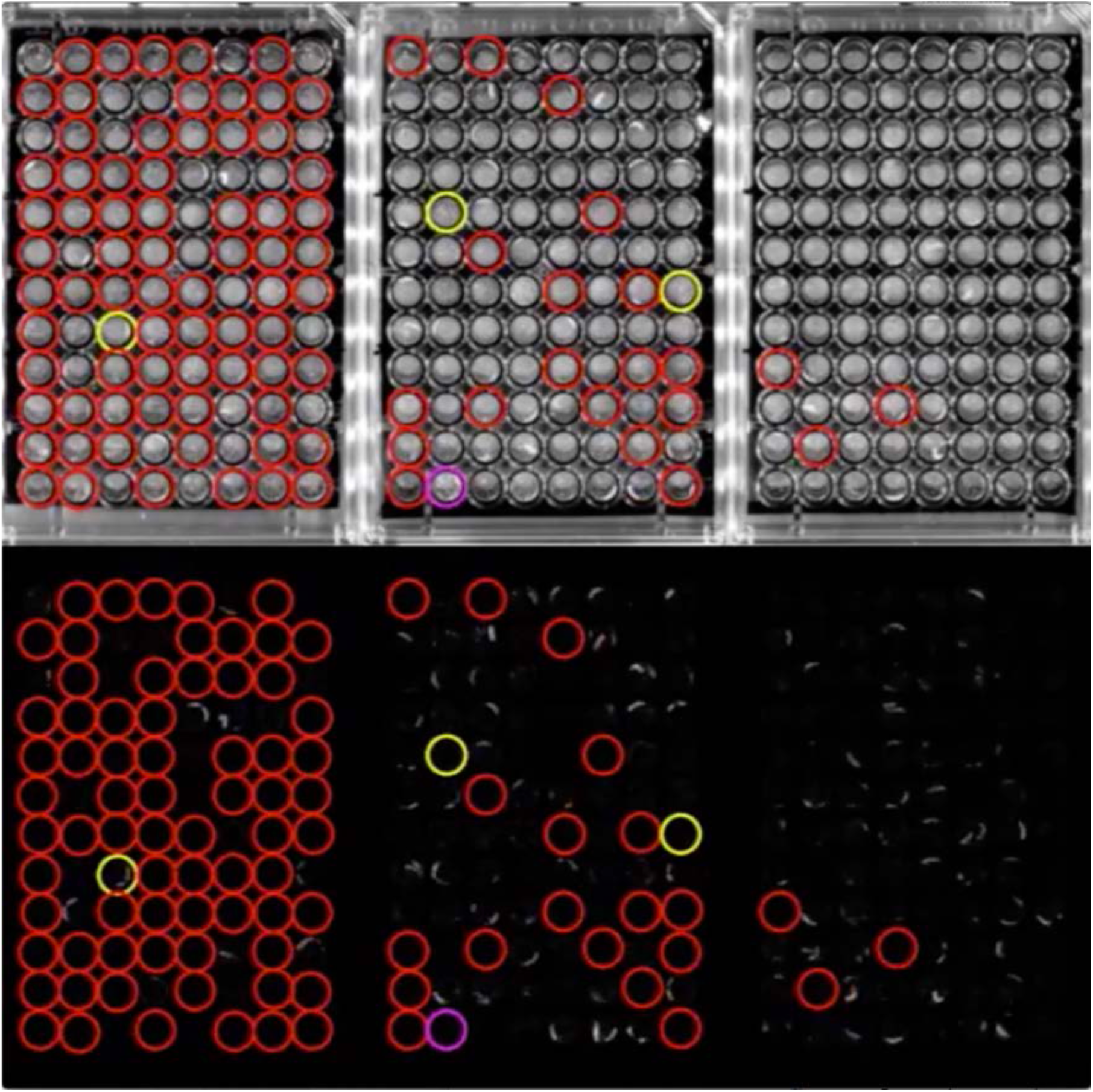
Dynamic visualization of life-event transition with single animal resolution in population image.

**Supplementary file 8A.**
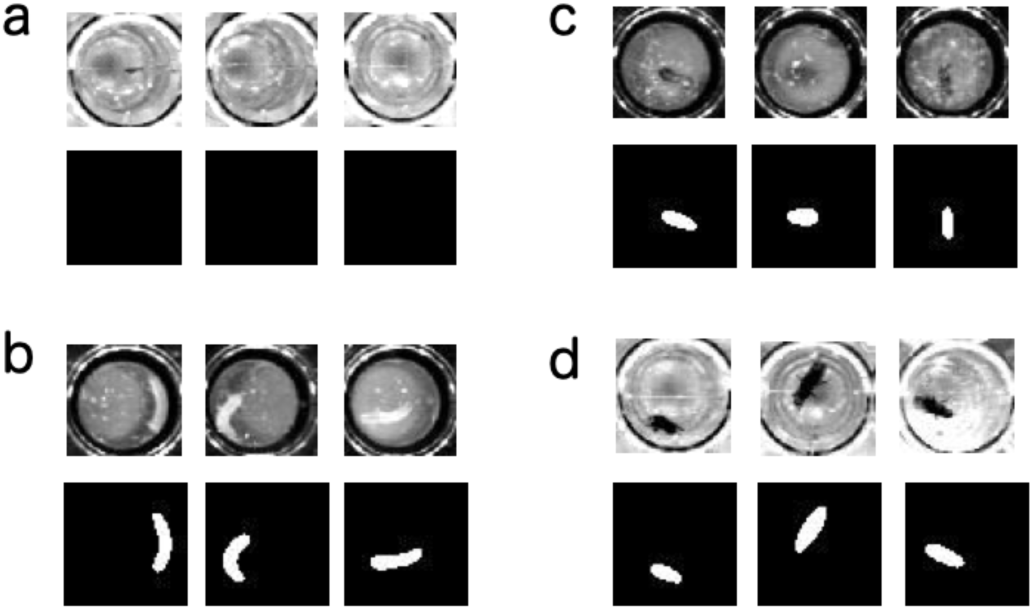
Example of segmentation for pupa (a), larva (b), adult in dataset 1 (c), and dataset 2 (d). Note that the segmentation correctly captured the animal body in the images obtained by different imaging condition (c and d), and correctly ‘ignored’ pupa (a).

**Supplementary file 8B.**
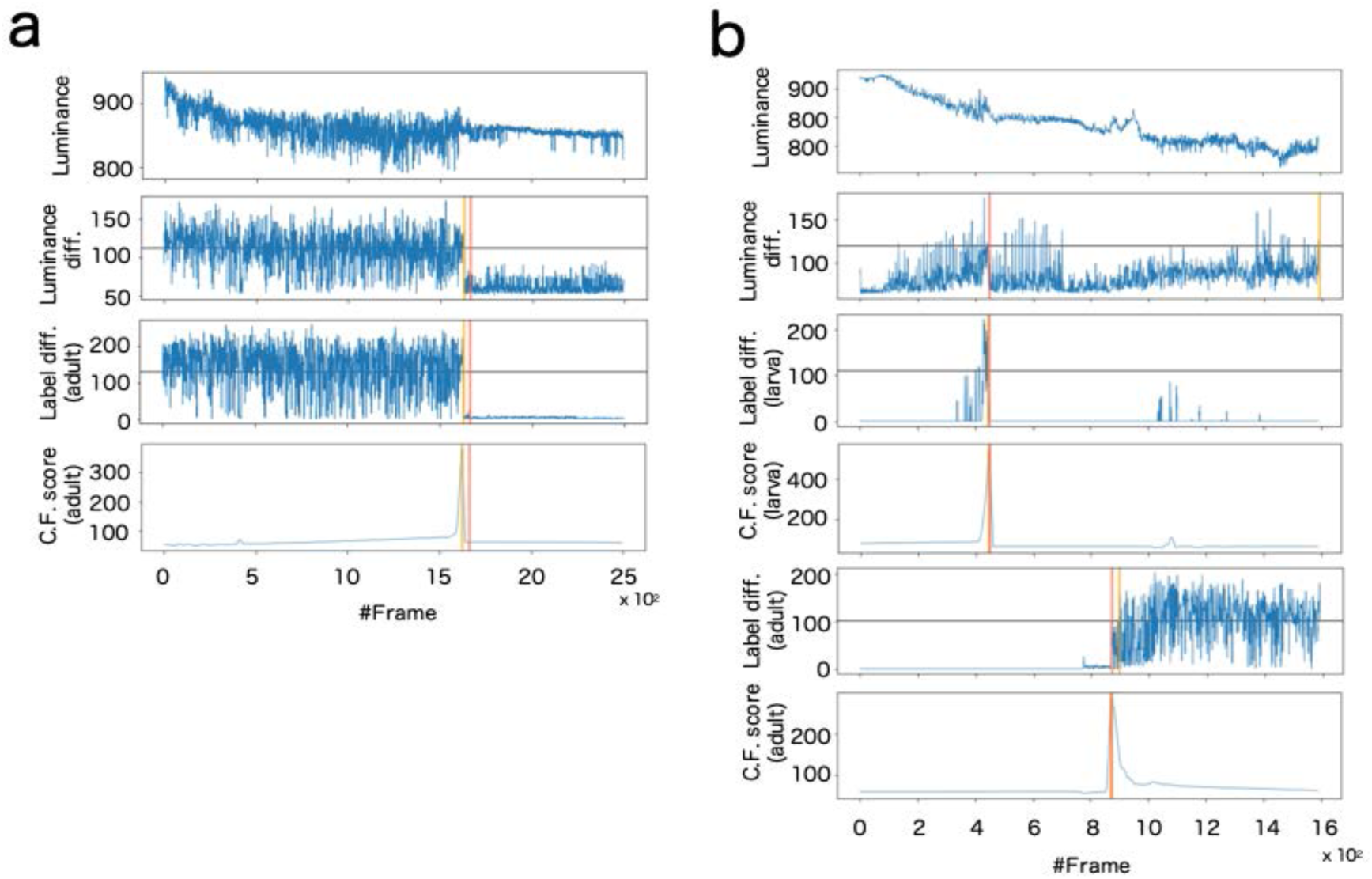
Example of signalization from image data and detection of death (a) and pupariation-eclosion (b). In (a), each panel indicates signals obtained as (*top*) luminance at every time steps, (*second panel*) subtraction of luminance between consecutive raw images, (*third panel*) subtraction between consecutive labeled images obtained by larva segmentation, and (*bottom*) ChangeFinder signal calculated by Label diff. Black horizontal lines indicate thresholds for detection of event timing. Red vertical line indicates the manually-detected event timings (pupariation), and yellow vertical lines are automatically-detected event timings (see *supplementary text*). In (b), almost same organization with (a) except both signals of larva and adult are included for detection of pupariation and eclosion.

